# Plasmid-driven strategies for clone success in *Escherichia coli*

**DOI:** 10.1101/2023.10.14.562336

**Authors:** Sergio Arredondo-Alonso, Anna K. Pöntinen, João Alves Gama, Rebecca A. Gladstone, Klaus Harms, Gerry Tonkin-Hill, Harry A. Thorpe, Gunnar S. Simonsen, Norwegian E. coli BSI Study Group, Ørjan Samuelsen, Pål J. Johnsen, Jukka Corander

**Author notes:** Equal contributions (Sergio Arredondo-Alonso, Anna K. Pöntinen, João Alves Gama, Rebecca A. Gladstone). Equal contributions (Ørjan Samuelsen, Pål J. Johnsen, Jukka Corander). Collaborators: Nina Handal, Nils Olav Hermansen, Anita Kanestrøm, Hege Elisabeth Larsen, Paul Christoffer Lindemann, Iren Høyland Löhr, Åshild Marvik, Einar Nilsen, Marcela Pino, Elisabeth Sirnes, Ståle Tofteland, Kyriakos Zaragkoulias (Norwegian E. coli BSI Study Group). Corresponding and ϕLead author: Jukka Corander.

## Abstract

*Escherichia coli* is the most widely studied microbe in history, but its extrachromosomal elements known as plasmids remain poorly delineated. Here we used long-read technology to high-resolution sequence the entire plasmidome and the corresponding host chromosomes from an unbiased longitudinal survey covering two decades and over 2,000 *E. coli* isolates. We find that some plasmids have persisted in lineages even for centuries, demonstrating strong plasmid-lineage associations. Our analysis provides a detailed map of recent vertical and horizontal evolutionary events involving plasmids with key antibiotic resistance, competition and virulence determinants. We present genomic evidence of both chromosomal and plasmid-driven success strategies that represent convergent phenotypic evolution in distant lineages, and use *in vitro* experiments to verify the importance of bacteriocin-producing plasmids for clone success. Our study has general implications for understanding plasmid biology and bacterial evolutionary strategies.

## Introduction

The extra-intestinal pathogenic *E. coli* (ExPEC) have been of particular interest to both eco-evolutionary and epidemiological research communities since they near-universally colonize the healthy human gut in an asymptomatic fashion, while also causing a high global burden of opportunistic infections, predominantly in the urinary tract and bloodstream^1^. ExPEC possess an arsenal of iron acquisition mechanisms, toxins and adhesins that enhance their persistence in this niche ^2–5^.

The ExPEC population is composed of a high diversity of clones, but a relatively small subset of lineages contributes disproportionately to the majority of urinary tract and bloodstream infections ^6^, where increasing levels of antibiotic resistance represent a challenge for effective treatment. Epidemiological investigations have uncovered an intriguing feature of ExPEC, showing stable maintenance of pandemic lineages over longer timescales and the emergence of novel successful clones in clinical surveillance, many with foodborne associations ^6–9^. The new clones typically expand rapidly initially, but are then constrained within only a few years such that the population transits to another equilibrium still maintaining a considerable diversity of lineages. Unbiased longitudinal genomic surveys have demonstrated that the transient disruption of the equilibrium bears the hallmarks of negative frequency-dependent selection (NFDS) ^10–12^, and further modeling work suggested this selection is acting on accessory genomic elements rather than on core genome variation ^13^.

Plasmids, initially described as transmissible agents carrying antibiotic resistance and competitor-inhibiting determinants in *E. coli* in the 1950s ^14–17^, play a key role in accessory genome dynamics. Numerous traits involved in antibiotic resistance, iron-acquisition mechanisms, bacterial competition, immune evasion and metabolism are encoded by genes residing on plasmids. In particular, some *E. coli* carry colicinogenic plasmids that produce bacteriocins: toxic proteins typically active against close relatives ^18^. Bacteriocins play an active role in intra-species competition ^19,20^ and have been proposed as one of the mechanisms maintaining NFDS ^21,22^. Despite the strong evidence that plasmids have played a critical role in the emergence and dissemination of successful clones ^23–26^, the structure of the plasmidome, and the horizontal and vertical evolutionary dynamics of plasmids and ExPEC hosts remain largely unknown. This is due to the paucity of scalable computational approaches to type complete plasmid sequences, and the lack of large-scale longitudinal long-read sequencing studies, which has hindered developing a firm understanding of the plasmidome structure and prevalence of particular plasmids.

Except in a recent study ^27^, ExPEC plasmids have previously been explored on a small-scale, most often in association with specific selective criteria, e.g. isolates carrying a particular resistance locus such as extended-spectrum □-lactamase (ESBL), and using mostly short-read sequencing data ^24,28–30^. To circumvent this, we long-read sequenced 2,045 *E. coli* isolates representing the genomic diversity of earlier short-read sequenced 3,245 bloodstream infection isolates from a 16-year national longitudinal study ^11^. These isolates were sampled regardless of their clonal background, antibiotic resistance profile or any other bacterial genotypic or phenotypic characteristic, making it ideal for the study of evolution, expansion and persistence of the plasmidome.

Our approaches resulted in a total of 4,485 circularized plasmid sequences and through inferred evolutionary rates and dated phylogenies we estimated the timing of plasmid emergence/acquisition into major ExPEC clones. We demonstrate multiple cases of plasmid-driven convergent evolution in distant genetic backgrounds, supporting the conclusion that plasmids play key roles in initial clonal establishment and later endemic maintenance in the host population. In particular, we found that some ExPEC clones were strongly associated with pColV-like plasmids encoding microcin V, a narrow-spectrum bacterial toxin involved in bacterial competition ^17^. By exploring the bacteriocin content present in the ExPEC population, we propose that plasmid-encoded bacteriocins play a pivotal role in shaping and maintaining the NFDS dynamics in *E. coli*. Through experimental approaches, we further demonstrate microcin V activity in different clonal backgrounds to inhibit the growth of multi-drug resistant ExPEC clones, thus contributing to the clonal success and maintenance of non-resistant clones in the population.

## Results

### Typing the ExPEC plasmidome: A network-based approach

To elucidate the ExPEC plasmid repertoire and its stability over time, we focused on a unique longitudinally sampled national epidemiological longitudinal collection (NORM collection) spanning a period of 16 years and involving 3,245 bloodstream infection isolates. This study included isolates regardless of their sequence type (ST) or antibiotic resistance background (e.g. presence of ESBL genes). To ensure the inclusion of isolates representative of the whole ExPEC population for long-read sequencing, we first selected 1,085 isolates (33.44%, 1,085/3,245) using a statistical method informed by the accessory genome diversity of the isolates (Fig. S1) based on short-read sequences ^31^. Next, we included all the remaining 960 isolates belonging to the four major STs, referred to as pandemic clones (ST69, ST73, ST95 and ST131) that caused nearly half of the bloodstream infections (44.19%, 1,438/3,254) (Gladstone et al. 2021). Thus, a total of 2,045 isolates (63,02% of the NORM collection) were selected for long-read sequencing, representing 216 distinct STs covering the entire ExPEC phylogeny (Fig. S2).

Highly contiguous assemblies were obtained for 1,999 genomes (Fig. S3A) with median N50 chromosome length 4.98 Mbp (Fig. S3B) and an average L50 chromosome count 1.09. A total of 5,417 contigs were identified as plasmid-derived, including 4,485 circular plasmid sequences, collectively here referred to as the plasmidome. Chromosome lengths differed significantly between the major clones (Fig. S3C, p-value < 0.05), where ST69 and ST73 showed a larger chromosome size on average compared to ST95 and ST131. The total length and the number of plasmid sequences observed per isolate was significantly lower for ST73 (p-value < 0.05) compared to the other major clones (Fig. S3D). This finding confirms previous reports showing that ST73 frequently carries lower plasmid load, demonstrated by lower conjugation frequencies and higher fitness costs ^32,33^.

To type the circular plasmid sequences (n = 4,485) identified from the hybrid assemblies, we first removed redundancy with cd-hit-est resulting in a set of 2,560 non-redundant plasmid sequences (Fig. S4). Next, we applied our previously developed method ‘mge-cluster’, that is well-suited for typing plasmid sequences derived from large studies ^34^. To ensure robust typing results, we clustered plasmids using a weighted undirected network based on multiple runs of mge-cluster with distinct parameter combinations. In the resulting network, nodes correspond to plasmids and edges to the association strength among plasmids co-clustering together in the distinct solutions (see Methods).

The network consisted of 15 components (connected subgraphs) encompassing 2,392 non-redundant plasmid sequences (93%) while 168 sequences remained as singletons (7%) (Fig. S4). Next, we used a popular community detection algorithm based on modularity optimization (Louvain method) to assign highly connected communities into non-overlapping groups (bins) ^35^. After quality control (see Methods), 2,285 non-redundant plasmids were clustered into 30 distinct groups referred to as plasmid types (pTs) labeled with two digits representing their network component and the community bin defined by the Louvain algorithm (Fig. S4). To obtain a consistent typing scheme and a compact visualization (Fig. 1), we selected the mge-cluster solution (perplexity = 62, cluster size = 49) closest (adjusted Rand index 0.99) to the network typing system.

**Figure 1.**
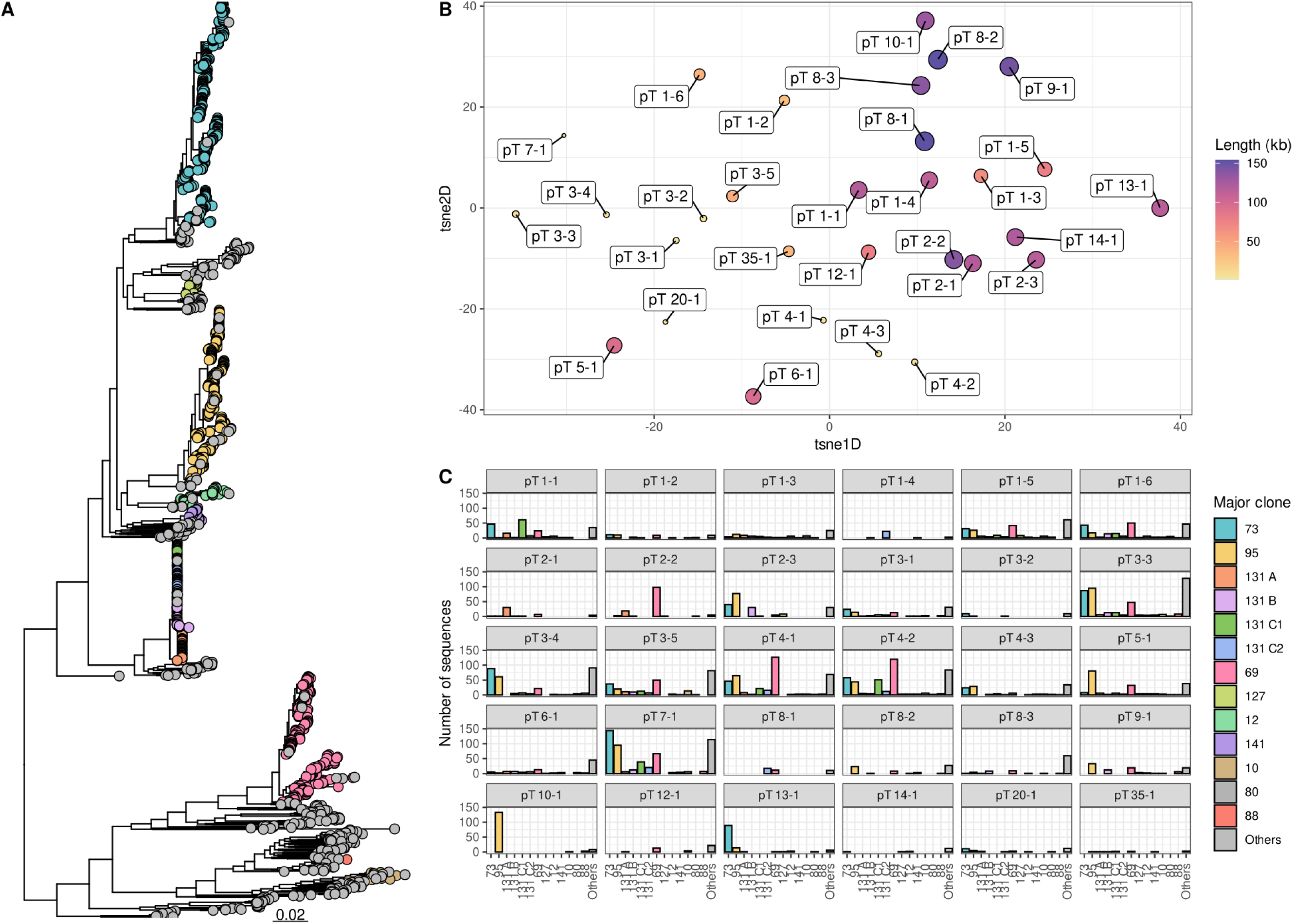
Plasmid diversity and typing scheme on the set of predicted plasmid sequences obtained from 1,999 ExPEC hybrid assemblies. A) Maximum-likelihood phylogeny of the 1,999 ExPEC isolates with an associated hybrid assembly. Isolates corresponding to the ten most prevalent sequence types (ST) in the NORM study are indicated with a distinct color and their position marked in the phylogeny. In the case of ST131, the distinct clades present in the clone (A, B, C1, C2) are represented. B) The closest mge-cluster solution (perplexity = 62, minimum cluster size = 49) to the network approach was considered to visualize the position of the pTs in the 2D embedding space. For visualization purposes, we computed the mean tsne1D and tsne2D coordinates of all sequences belonging to each pT and considered those coordinates to represent the position and anchor each pT in the embedding. Each pT is labeled with its associated pT consisting of two digits indicating their graph component and associated bin and its size is proportional to the average plasmid length. C) Barplot indicating the total number of plasmid sequences assigned to each pT and belonging to the main STs found in the ExPEC collection

Finally, the redundant sequences (n=1,925) discarded by cd-hit were assigned to the same pT as their representative sequence (Fig. S4). Using the existing operational mode of mge-cluster, we could further assign 498 non-circular plasmid sequences to specific pTs without the need to re-consider the entire network and to modify the typing scheme. In total, 4,569 plasmid sequences (84%, 4,569/5,417: 4,071 circular and 498 non-circular) were assigned to the 30 defined pTs (Fig. S4) representing the ExPEC plasmidome.

### High levels of gene sharing and genetic relatedness of IncF plasmids

To link our assigned pTs with existing plasmid classification and typing schemes, we identified the dominating replicons (PlasmidFinder), plasmid MLST schemes (pmlst) and checked their congruence with another well-established plasmid typing tool (MOB-suite) (Table S2). In a previous publication ^34^, we showed that mge-cluster and MOB-suite were concordant on the typing of *E. coli* plasmids, but mge-cluster was generally better at grouping together sequences with a shared plasmid backbone. For each pT, we further determined the degree of gene sharing, predicted mobility (conjugative, mobilisable or non-mobilisable) and the presence of antibiotic and virulence genes (Table S2).

Our analyses revealed that all pTs with a large average plasmid size (> 100 kb) contained IncF variant replicons, harbored antibiotic and/or virulence determinants, and displayed high degrees of gene sharing (Fig. S5) as measured by their containment index ^36^. Our results support large plasmids as a key resource of bacterial adaptive traits. In contrast, pTs belonging to small sequences (< 10kb) only shared a minor fraction of their genome content corresponding to the genes involved in their replication or mobilization machinery (Fig. S5).

Gene synteny analysis (minimum identity 0.8) among representative sequences of large IncF pTs (> 100 kb) (Fig. 2) revealed the presence of large genomic blocks in common including siderophore-operons (*iroBCDEN*, *iutA*-*iucABCD* and *sitABCD* operons), antibiotic resistance genes and the prototypical conjugation F-type transfer system. By increasing the stringency to create a link between genes in synteny (minimum identity 0.99) (Fig. S6), we observed that the F-type transfer region diversified overall more than other genomic blocks. Thus, our data strongly suggests that pTs 8-2/9-1/10-1 and pTs 2-1/2-2/2-3 have recently evolved from the same ancestral pT respectively, since their transfer regions remain highly conserved and they differ mainly by the acquisition of antibiotic resistance genes. We confirmed this by reconstructing a recombination-free phylogeny using the aligned plasmid sequences from each combination of plasmid types (Fig. S7), which showed that pT 8-2 and pT 2-3 were the ancestral plasmid types. Furthermore, pT 9-1 and pT 2-2 arose respectively from pT 10-1 and pT 2-1. In addition, these pTs were clustered in the same group by MOB-suite confirming their relatedness (Table S2).

**Figure 2.**
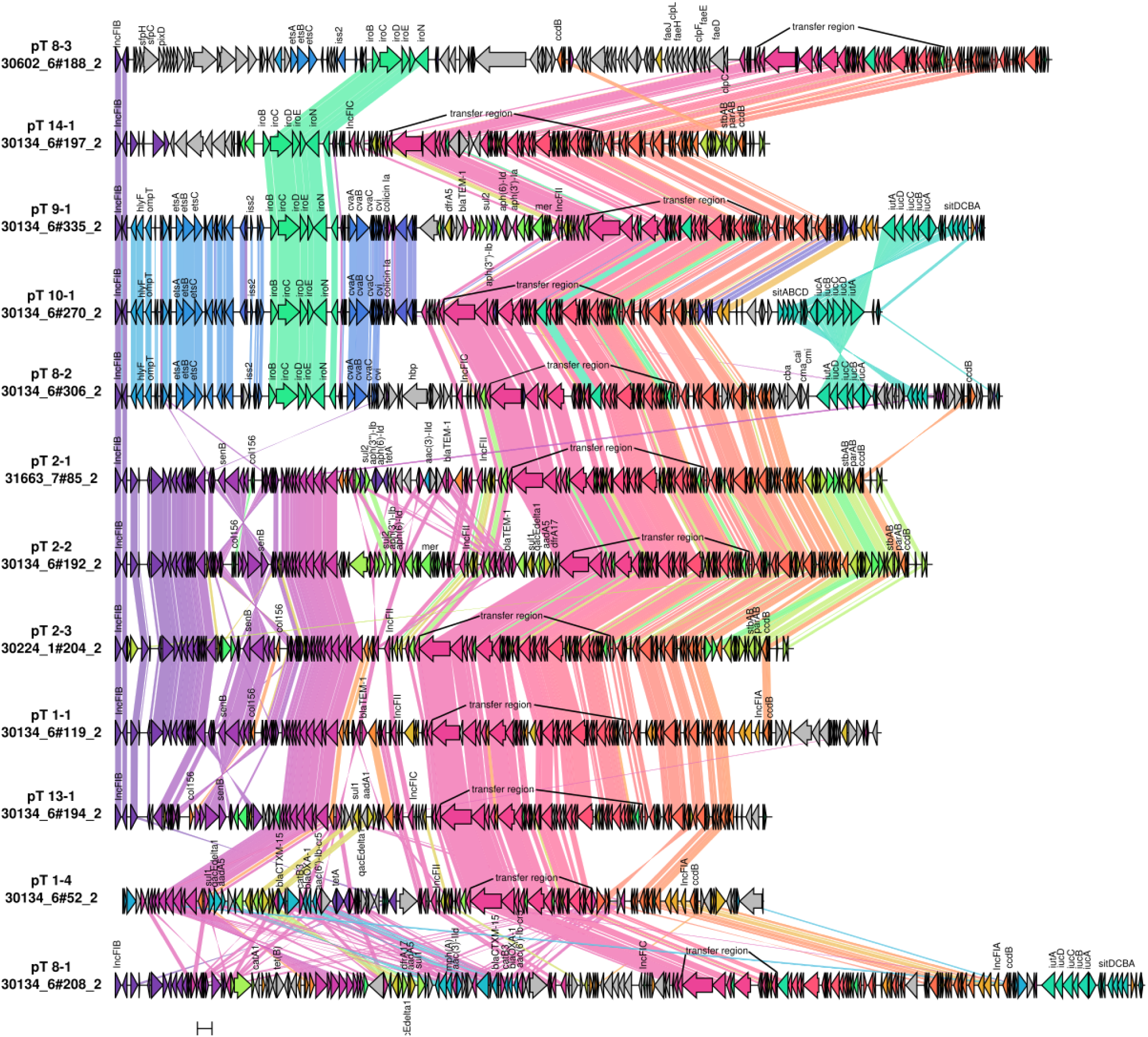
Synteny analysis and visualization of the main IncF large plasmid types (pT). For each pT, we selected a plasmid sequence (tagged with its sequence ID) representing the most predominant features present in the cluster (as summarized in Table S2). The gene synteny analysis was performed using clinker ^88^ considering a minimum identity threshold of 0.8 to draw a link between genes. Virulence and antibiotic resistance genes were identified using the EcOH database ^82^ (indexed in Abricate) and AMRFinderPlus ^83^ and their names curated according to well-studied ExPEC reference plasmids.

### Unraveling the shared evolutionary history of plasmid and clones

After elucidating and typing the ExPEC plasmidome, we explored whether particular pTs could have contributed to the success of pandemic ExPEC clones. We focused our analyses on pTs corresponding to medium and large plasmids (> 10kb) and in more detail on pColV-like and pUTI89-like plasmids which have previously been linked to: i) ExPEC virulence, ii) adaptive traits influencing host range, iii) zoonotic potential, and iv) antibiotic resistance carriage ^37–40^. To this end, we inferred the evolutionary rates and dated the phylogenies of the four major clones (ST69, ST73, ST95, ST131) using BactDating ^41^ and further detected probable clonal expansions associated with the acquisition of pTs using CaveDive ^42^. This permitted us to estimate when ExPEC clones acquired specific pTs and whether a clonal expansion was followed by that association.

ST95, an ExPEC clone mostly susceptible to antibiotics, showed a high prevalence (37%) of the pColV-like plasmid pT 10-1 (Fig. S8). As illustrated in Figure 2, this plasmid possesses all features representative of pColV-like plasmids ^43^ including several iron-acquisition mechanisms (salmochelin, aerobactin), bacteriocin genes involved in clone competition (microcin V, colicin Ia)^38,44^, immune evasion genes conferring resistance to serum (*iss) ^39^* and virulence genes (e.g. *hlyF) ^45^*. The distribution of pT 10-1 in the phylogeny of ST95 (Fig. 3A) suggests that the plasmid was introduced by the most recent common ancestor (MRCA) of ST95 dating back to 1768 (95% CI, 1721–1806). Furthermore, the major branch leading from the root of ST95 was identified as a clonal expansion occurring in 1823 (95% CI, 1789-1851). By comparing plasmid versus core-genome distances, we observed a positive correlation (0.78) between pT 10-1 and ST95 phylogenies further supporting that the plasmid has been vertically transmitted in ST95 instead of being repeatedly acquired (Fig. S9A). In some instances, we observed the replacement of pT 10-1 with pT 9-1, the alternate version of the pColV-like plasmid characterized by the presence of an antibiotic and heavy-metal resistance gene cassette. A particular clade of ST95 (fimH41:O1/O2:H7) showed multiple and independent replacements of pT 10-1 with pT 2-3 (Fig. S8, Fig. S9B), a pUTI89-like plasmid labeled by the pmlst scheme as F29:A-:B10. This plasmid has been characterized by its immune evasion properties ^46^, and by playing an important role in virulence and biofilm regulation (*senB and parAB* genes) ^47,48^.

**Figure 3.**
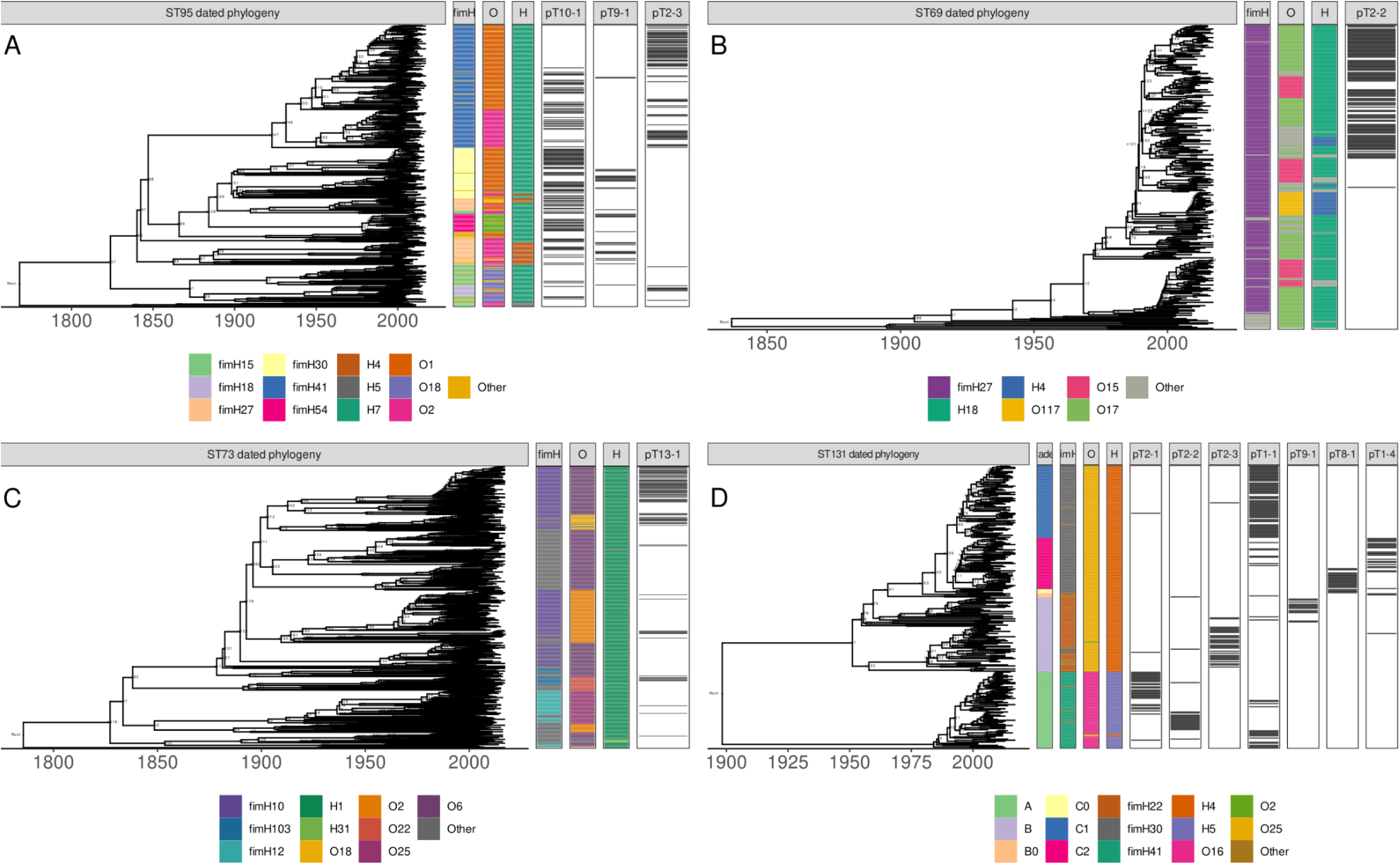
Dated phylogenies of the four main ExPEC clones and presence/absence pattern of the most prevalent plasmid types (pT) found in each sequence type (ST). A) Dated phylogeny of ST95 with clades indicated based on fimH/O/H typing and presence/absence of the pcolV-like plasmids (pTs 10-1 and 9-1) and the pUTI89-like plasmid (pT 2-3). B) Dated phylogeny of ST69 with clades indicated based on fimH/O/H typing and presence/absence of the pUTI89-like plasmid (pT 2-2). C) Dated phylogeny of ST73 with clades indicated based on fimH/O/H typing and presence/absence of the pT 13-1. D) Dated phylogeny of ST131 with clades indicated based on existing nomenclature (A, B, B0, C0, C1, C2), fimH/O/H typing and presence/absence of multiple associated pTs.

The basal part of the ST69 phylogeny represents isolates notably void of larger plasmid carriage, suggesting that such plasmids did not play a central role in the early evolution of this lineage (Fig. 3B). However, a particular clade of ST69 (fimH27:017/015:H18/H4) showed the ubiquitous presence of the plasmid type pT 2-2, a pUTI89-like plasmid version highly similar to the pT 2-3 described in the specific clade of ST95. In contrast, this plasmid harbors several antibiotic resistance genes providing resistance to multiple antibiotic groups (e.g. β-lactams, aminoglycosides, sulphonamides and trimethoprim) and to heavy-metals (mercury) (Fig. 2). The plasmid type was estimated to be acquired around ∼1990 (95% CI, 1986-1992) and notably this acquisition was nested with a major expansion event occurring two nodes earlier in the tree (1987, 95% CI, 1984-1990).

Consistent with its generally smaller inferred plasmidome size (Fig. S3D), ST73 only carries large plasmids in a minority of isolates (181/560, 32%). Out of these, the plasmid type pT 13-1 (prevalence 18%) (Fig. S8) encoding similar virulence traits as pUTI89-like plasmids (e.g. *senB*) has been introduced multiple times into ST73. In particular, we detected a clonal expansion event in 1983 (95% CI, 1977-1989) after the plasmid had acquired the β-lactamase *bla*_SHV-1_ gene, displaying increased ampicillin resistance, around 80 years after the initial pT 13-1 acquisition (1900, 95% CI 1877-1919) into the ST73 fimH10:O6:H1 clade (Table S3, Fig. 3C). This IncF plasmid (pmlst F51:A-:B10) frequently encoded several antibiotic resistance genes including β-lactam (*bla*_SHV-1_, *bla*_TEM-1_), aminoglycoside (*aadA1*, *aph(3’’)-Ib*, *aph(6)-Id*) and sulphonamide (*sul1*, *sul2*) resistance genes (Table S1) and has been shown to be associated with a higher carriage of resistance genes within ST73 ^49^. Despite the general trend of susceptibility for this clone, this demonstrates that particular subclades of ST73 can become multi-drug resistant by incorporating this plasmid type.

In agreement with previous reports based on short-read sequencing data ^24,28^, we observed strong plasmid-clade associations between particular IncF pTs and ST131 clades. In clade A (fimH41:016:H5), the most basal clade of ST131, pT 1-1 was present in the MRCA of the clade (1984, 95% CI 1978-1989) and later replaced around 1991 (95% CI, 1986-1996) by pUTI89-like plasmids (pTs 2-1, 2-2). The branch where this plasmid type replacement occurred was inferred to correspond to a clonal expansion event estimated to have occurred around 1992 (95% CI, 1988-1996), recapitulating the detected association between plasmid and clonal expansion observed for ST69.

In clade B (fimH22:025:H4), which is generally strongly associated with antibiotic susceptibility ^11^, we observed a similar plasmid distribution as described above for ST95. The pColV-like plasmid, pT 9-1, was introduced in a sublineage of clade B around 1955 (95% CI, 1944-1964) while the pUTI89-like plasmid (pT 2-3) entered between 1957 and 1981 (95% CI, 1946-1986) in another clade B sublineage (Fig. 3D, Table S3).

Finally, we explored the plasmid associations in clade C (fimH30:025:H4) which is characterised by *gyrA* and *parC* mutations conferring fluoroquinolone resistance, and further subdivided into C1 (associated with the *bla*_CTX-M-27_ allele) and C2 lineages (associated with *bla*_CTX-M-15_). ST131-C1 showed a high prevalence of pT 1-1 (pmlst F1:A2:B20) (Fig. S8). Its distribution suggests that this plasmid was present in the MRCA of ST131-C1 (∼1993, 95 CI 1989-1996) (Fig. 3D, Table S3). This plasmid encodes for the *senB* gene cluster and frequently harbours multiple non-fixed antibiotic resistance genes, including *bla_TEM-1_*, *bla_CTX-M-27_* genes (β-lactams), or *aac(3)-IId* (aminoglycoside) among others (Table 1). In contrast, ST131-C2 harboured pT 8-1 (pmlst F36:A4:B58) (Fig. 2) which was also present in its basal subclade (C0) (Fig. 3D). This plasmid type contained a variable set of antibiotic resistance genes including the *bla*_CTX-M-15_ ESBL gene, *dfrA17* (trimethoprim), *aadA5* (aminoglycoside), and *sul1* (sulfonamide) (Fig. 2, Table S1). pT 1-4 (pmlst F2:A1:B- Table S2) was introduced later in the ST131-C2 clade (∼1992, 95%CI 1988-1996), rapidly increasing in prevalence and including the *bla*_CTX-M-15_ ESBL gene.

Taken together, IncF pColV- and pUTI89-like plasmids played a key role in the evolution of pandemic ExPEC clones and are associated with clonal expansion events often linked to antibiotic resistance gene acquisition.

### Plasmid-encoded bacteriocins play a pivotal role in bacterial competition

We previously showed that antibiotic resistance is not the sole driver of clonal success, highlighted in particular by the stable circulation of ST95, ST73 and ST131-B clades which all remain largely susceptible to most classes of antibiotics ^11^. Our data suggest the presence of bacteriocins encoded in pColV-like plasmids for both ST95 and ST131-B could play a fundamental role in bacterial competition. To investigate the role of bacteriocins across the entire set of lineages, we expanded our analysis by searching for genes coding for bacteriocins regardless of their genomic location.

The majority of identified bacteriocin genes were located on plasmids (Fig. S10A), with the exception of *mchB*, encoding microcin H47, typically found in the chromosome of ST73 isolates (Fig. S10B). The most frequently found plasmid-encoded bacteriocins were *cvaC* (microcin V) and colicin Ia (also termed as pECS88_0104) (Fig. S10A), typically co-occurring in the pColV-like plasmids (Fig. 2, Fig. S10A) and associated with ST131-B and ST95, respectively (Fig. S10B). Some of the pTs with smaller average sizes (pTs 3-3, 3-4) occasionally carried bacteriocins (Fig. S10A) and were found in a wide diversity of STs indicating multiple acquisitions/losses of the genes, but showed no evidence of clonal expansions in contrast with pColV-like plasmids (Fig. S10B).

To elucidate the role of microcin V and colicin Ia in clone competition, we examined their activity experimentally. The microcin V gene cluster is composed of four genes (Fig. 4A): *cvaC* encodes the toxin, the immunity gene *cvi* encodes a peptide that protects the toxin-producing cells, and the genes *cvaA* and *cvaB* encode a secretion system to secret the toxin ^17^. Production of microcin V is induced by iron limitation, and in target cells recognized by the siderophore receptor Cir ^17^. When grown in iron-limiting conditions, the culture supernatants of three ST95 and one ST131-B plasmid-harboring isolates strongly inhibited the growth of the *E. coli* lab strain MG1655 (Fig. 4B). The killing effect was dose-dependent, which is consistent with bacteriocin activity and the absence of visible plaques ruled out bacteriophage-mediated killing. In the absence of induction, the ST95 isolate 27-61 (Table S4) produced a weaker inhibition zone, possibly due to some basal expression of one of the two bacteriocins. As a further control, the ST131-B isolate 27-56 not harboring a bacteriocinogenic plasmid was unable to produce inhibition despite growing under iron-limiting conditions (Fig. 4B). The isolates 31-16 and 31-17 (Fig. 4) served as further controls (see Methods).

**Figure 4.**
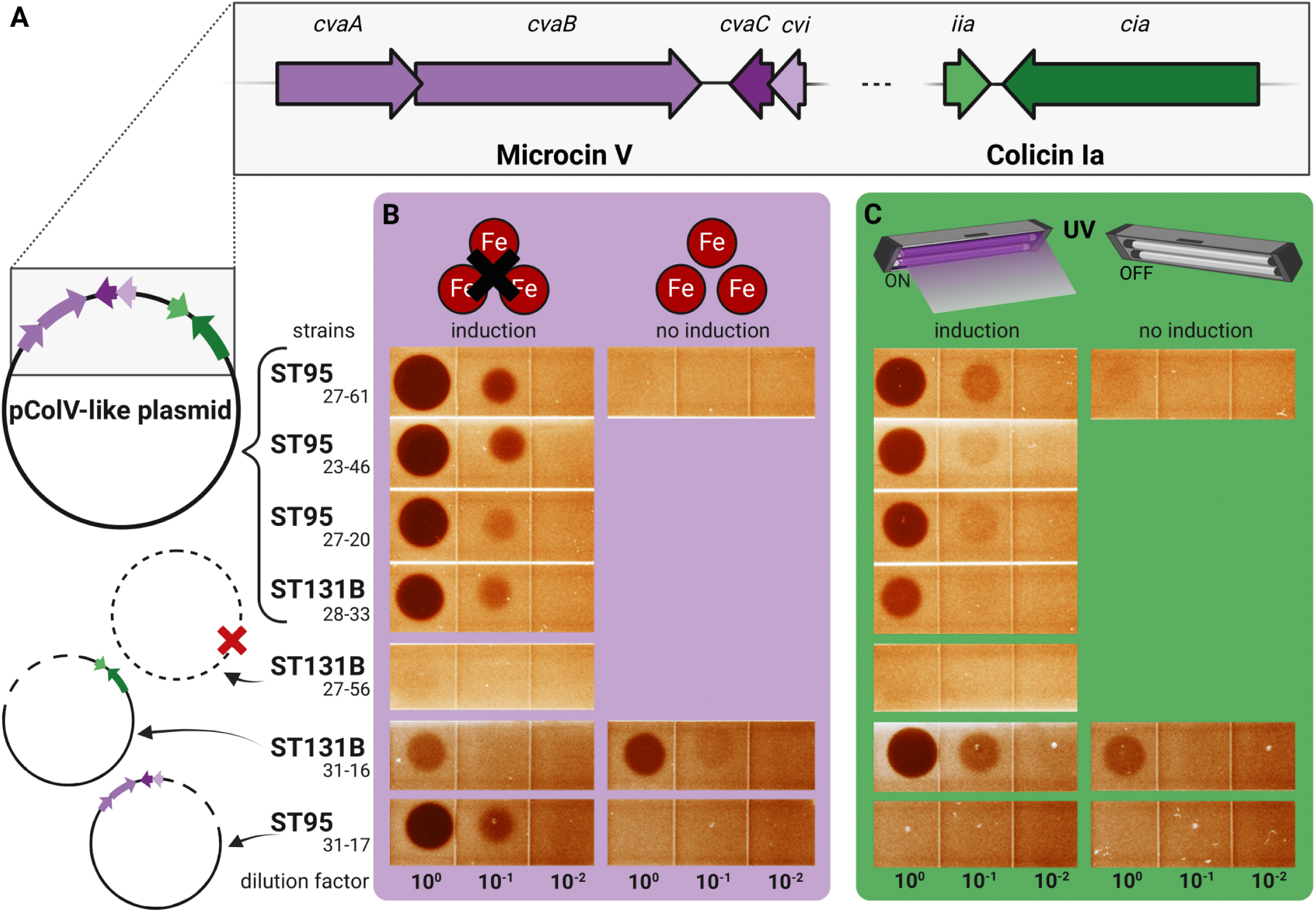
Schematic representation of the bacteriocin susceptibility assay. A) Description of bacteriocinogenic plasmids from pT 9-1 (ST131 clade B) and pT 10-1 (ST95). These archetypical pcolV-like plasmids encode two bacteriocin gene clusters: microcin V and colicin Ia. Darker and lighter arrows indicate respectively the toxin (*cvaC, cia*) and immunity genes (*cvi, iia*) of each cluster. Control isolates either harbor no plasmid (ST131 clade B isolate 27-56) or harbor plasmid versions encoding only a single bacteriocin (colicin Ia for ST131 B isolate 31-16, microcin V for ST95 isolate 31-17). B) Susceptibility of *E. coli* MG1655 to microcin V. Production of microcin V is induced by iron limitation (simulated experimentally with the addition of the chelating agent 2,2’-bipyridyl). Bacteriocinogenic activity is demonstrated by growth inhibition in the spotted area (in a dilution-dependent way and without formation of individual plaques). C) Susceptibility of *E. coli* MG1655 to colicin Ia. Production of colicin Ia is induced through SOS induction (stimulated experimentally with UV irradiation). Bacteriocinogenic activity is demonstrated as in B. This figure was created with BioRender.com

Having demonstrated that microcin V is functional, we further tested its range of activity against 51 *E. coli* isolates belonging to the four major clones STs, as well as against ST10, which is an ancestral clone ubiquitously present in the gut of mammals and avian species. As expected, the bacteriocin-producing isolates mentioned above all displayed resilience to any of the corresponding four batches of microcin V, while 39 of the remaining 47 isolates were sensitive (Fig. S11, Table S4). Sensitive isolates were generally inhibited by all four batches, except in a few cases where inhibition was only observed for the more concentrated bacteriocin batches (produced by isolates 23-46 and 27-61). Overall, ST69 and ST131 isolates were sensitive to the action of microcin V, while isolates belonging to the other STs display a more heterogeneous behavior. The supernatants of uninduced 27-61 and induced 27-56 (lacking colicinogenic plasmid) did not affect growth on the clinical isolates, as expected.

The colicin Ia gene cluster is composed of two genes (Fig. 4A): *ciaA* encoding the respective toxin, and *iia* that encodes immunity. This colicin also uses the Cir receptor, and is induced by the SOS response ^44^. The same four plasmid-harboring isolates used above were exposed to UV irradiation to induce the SOS response and consequently production of colicin Ia. All four culture supernatants inhibited the growth of *E. coli* MG1655 in a dose-dependent manner with no observable phage plaques (Fig. 4C). As shown for microcin V, the uninduced ST95 isolate 27-61 produced a weaker inhibition zone, while the induced ST131-B isolate 27-56 not harboring a colicinogenic plasmid was not affected.

Testing the four supernatants against the 51 isolates mentioned above, only six displayed sensitivity (Fig. S11, Table S4). Supernatants of isolates 27-56 (induced, no plasmid) and 27-61 (uninduced) did not display inhibitory activities

Altogether, these results show that pColV-like plasmids present in ST95 and ST131-B encode functional bacteriocins, and in particular we show the microcin V gene cluster inhibits the growth of a wide range of *E. coli* isolates from different genetic backgrounds supporting their role in clone competition and possibly NFDS maintenance.

## Discussion

Plasmids are generally considered as the ultimate vehicle of rapid evolution in bacterial genomes, while also sometimes regarded as parasitic elements to their host cells ^14–16,50^. Our analysis indicated that larger plasmids in *E. coli* are almost always associated with traits falling under bacterial competition, pathogenesis and antibiotic resistance. Even many of the small plasmids were found to incorporate likely beneficial bacteriocin-producing systems, and were scattered around the population phylogeny, indicating their repeated horizontal acquisition, but lacking evidence of further clonal expansion.

Only two co-occurring plasmid-encoded bacteriocins (microcin V and colicin Ia) were frequent enough and showed evidence of clonal expansion in different genetic backgrounds to indicate a major role in colonization competition and contribution towards maintaining stable population equilibrium. To further examine their role in competition, we experimentally confirmed the ability of microcin V to inhibit the majority of frequently observed clones in the population that lack this system. Notably, this bacteriocin shows a stable association with lineages/clades displaying a notable degree of susceptibility to antibiotics (ST95, ST131-B) and is able to inhibit isolates from the most successful multi-drug resistant lineages/clades (ST69, ST131-C1, ST131-C2), suggesting it is an important factor contributing to the continued maintenance of the susceptible lineages in the population. Similarly, only a single dominant chromosomal bacteriocin, the microcin H47, was identified in our collection. Characterics of these bacteriocins thus stand in stark contrast with the substantial bacteriocin diversity previously found to contribute to the maintenance of a stable equilibrium in the *Streptococcus pneumoniae* population over time ^22^. However, two ST10 isolates were found insensitive to microcin V, despite lacking this bacteriocin-producing system. This is well aligned with the frequently observed colonization ability of ST10 and its lower virulence potential ^51^ that was recently quantified at the population level by systematically comparing neonatal colonization rates with those observed in bloodstream infections ^5^. Why a subset of ST10 population would remain insensitive to microcin V is currently unclear and warrants further investigation into evasion of the effects of these inhibition mechanisms.

The unique characteristics of our dataset allowed detection of multiple events of convergent phenotypic evolution across distant lineages, associated with acquisition and stable maintenance of certain plasmid types over time. In addition to the same plasmid-based bacteriocin-producing systems observed in ST95 and ST131-B discussed above, these two distant lineages display congruent phenotypes also in other clades, that host plasmids different from the pColV-like plasmids. These correspond to the widely studied reference plasmid pUTI89, encoding virulence traits archetypal for uropathogenesis ^46^. In both evolutionary backgrounds, these clades were non-overlapping with the clades hosting pColV-like plasmid, and have been found to be strongly associated with avian pathogenic *E. coli* ^25,38,52^. ST131-B has been previously widely indicated as a foodborne uropathogen, in particular related to consumption of poultry meat ^53,54^. Combined, this evidence indicates that ST95 and ST131-B lineages circulating in human hosts are stably divided into human vs. avian adapted subclades assisted by their plasmid-associated traits, and that the latter likely reflects a constant spillover from the avian niche via foodborne transmission. Interestingly, the estimated timing of the pColV-like plasmid acquisition in ST131-B during 1950-60s coincides with a rapidly intensifying poultry production in many countries after WW2, which may have provided an opportunity for *E. coli* equipped with particular plasmid-derived traits to quickly expand in this niche.

Similar to the previously discussed observations, ST69 and ST131-A display clades characterized by a shared plasmid content, which we showed arose from the pUTI89-like virulence plasmids and absorbed additional resistance genes that remain in a stable association with the virulence determinants. This exemplifies plasmid-associated convergent phenotypic evolution in two lineages belonging to different phylogroups (D, B2). ST69 was originally detected in an outbreak in California ^55^, and later epidemiological investigation suggested it emerged in the late 1990s and then became endemic globally within 10 years ^56^. Our dating analysis gives the narrow time interval between 1987-1992 for the acquisition of the plasmid which ultimately led to a global dissemination of this clone with the specific virulence and antibiotic resistance characteristics. Further, as shown by our results, this evolutionary process closely reflects the parallel acquisition of the same plasmid by ST131-A subclade, which is again associated with remarkably similar epidemiological characteristics. i.e. a rapid expansion ^11,12^ and global endemicity over time. These two evolutionary trajectories across the successful lineages of *E. coli* bear the same hallmarks as the convergence of hypervirulence and multi-drug resistance in *Klebsiella pneumoniae*, which were first identified as non-overlapping traits ^57^, but later studies pinpointed to plasmid-driven merging of the two in particular genetic backgrounds ^58,59^.

Our observations of multiple cases of convergent evolution in distant genetic backgrounds by plasmids as a vehicle, suggests that they function as a significant means for new clones to first establish, and then later endemically sustain themselves in a host population. Future studies, including mechanistic modeling of plasmid evolution and maintenance, could shed further light on the factors that clone success ultimately depends on.

### Limitations of the study

Despite the strong association between specific plasmid types, we observed that some isolates across the phylogeny have lost their associated plasmid. The normalized coverage of those plasmid types in the hybrid assemblies suggests that a proportion of the cells sequenced have lost the plasmid as they are present in a copy number lower than the chromosome. Such lower coverage could be explained by deletions of large plasmid regions, which can be adaptive by rendering plasmid variants less costly relative to the original plasmid ^23,60^. Plasmid stability with its bacterial host is dependent on the ecological context, and some of the isolates could lose the associated plasmid when cultivated in the lab environment ^61^. In addition, some *E. coli* plasmids carrying ESBL genes have lower fitness costs for the population when present in low frequencies ^62^. Altogether, these factors could thus hide the true prevalence of some of the plasmid types we identified in the ExPEC clones. It should further be noted that the current analysis does not prove the causal roles of the plasmids that were found to be associated with the inferred clonal expansions, despite that the combined evidence from functional characterization and population genomic modeling is strong. Indeed, predicting the stability of a plasmid in a host population is a difficult endeavor ^63–66^. Plasmid-host associations result from multiple factors such as the level of benefit provided by the plasmid ^67^, the ability to acquire compensatory events during co-evolution ^65^, the pre-existence of host genotypes that favor the acquisition/maintenance of a plasmid ^67–70^, as well as the ecological context ^71,72^. Definite causal quantification of the impact of these plasmids is challenging to obtain since the conditions for their expansions in the human host population are very difficult to replicate in any experimental setting and a counterfactual analysis remains practically an impossibility in this ecological context.

## STAR Methods

### Key Resources Table

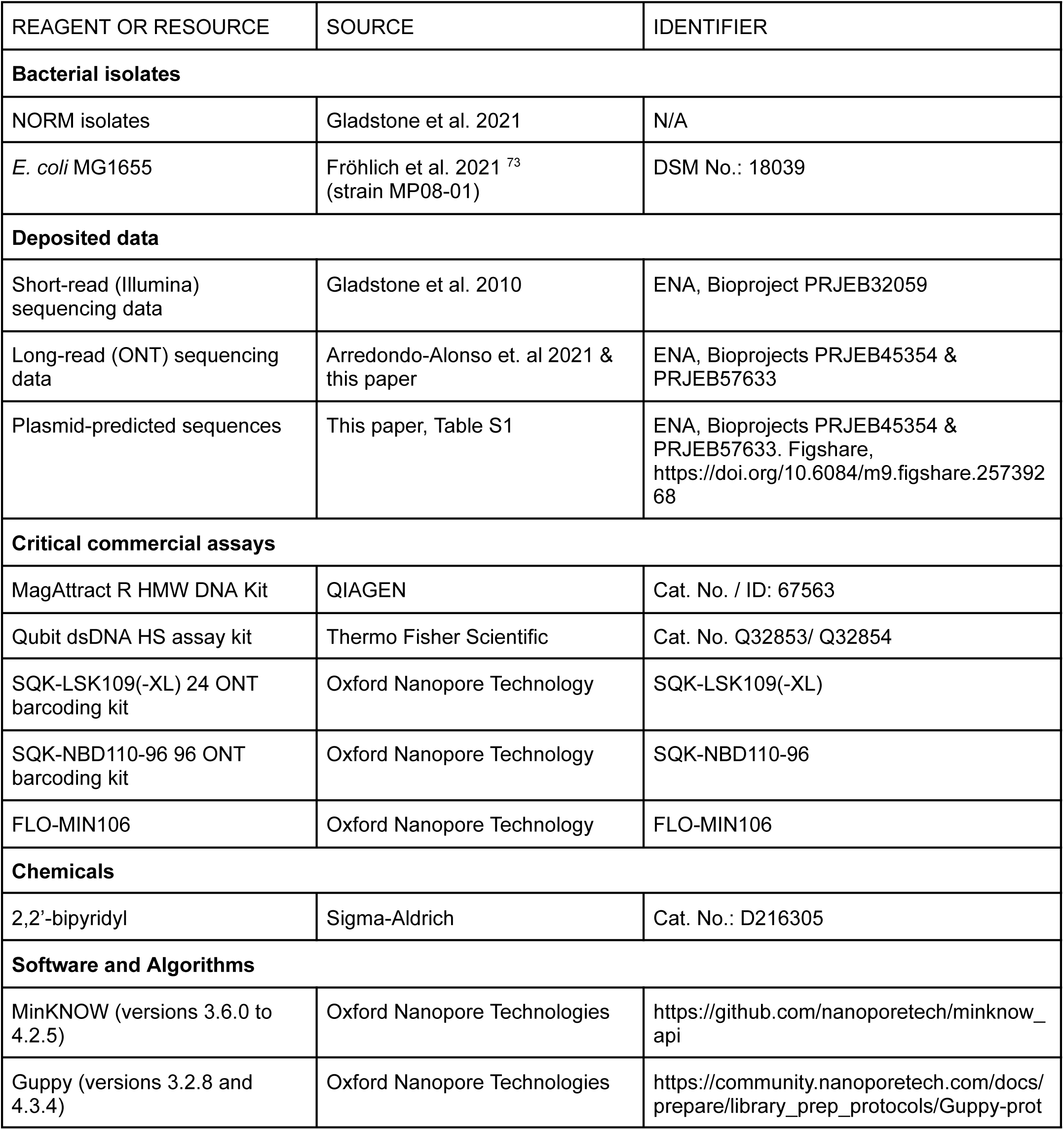
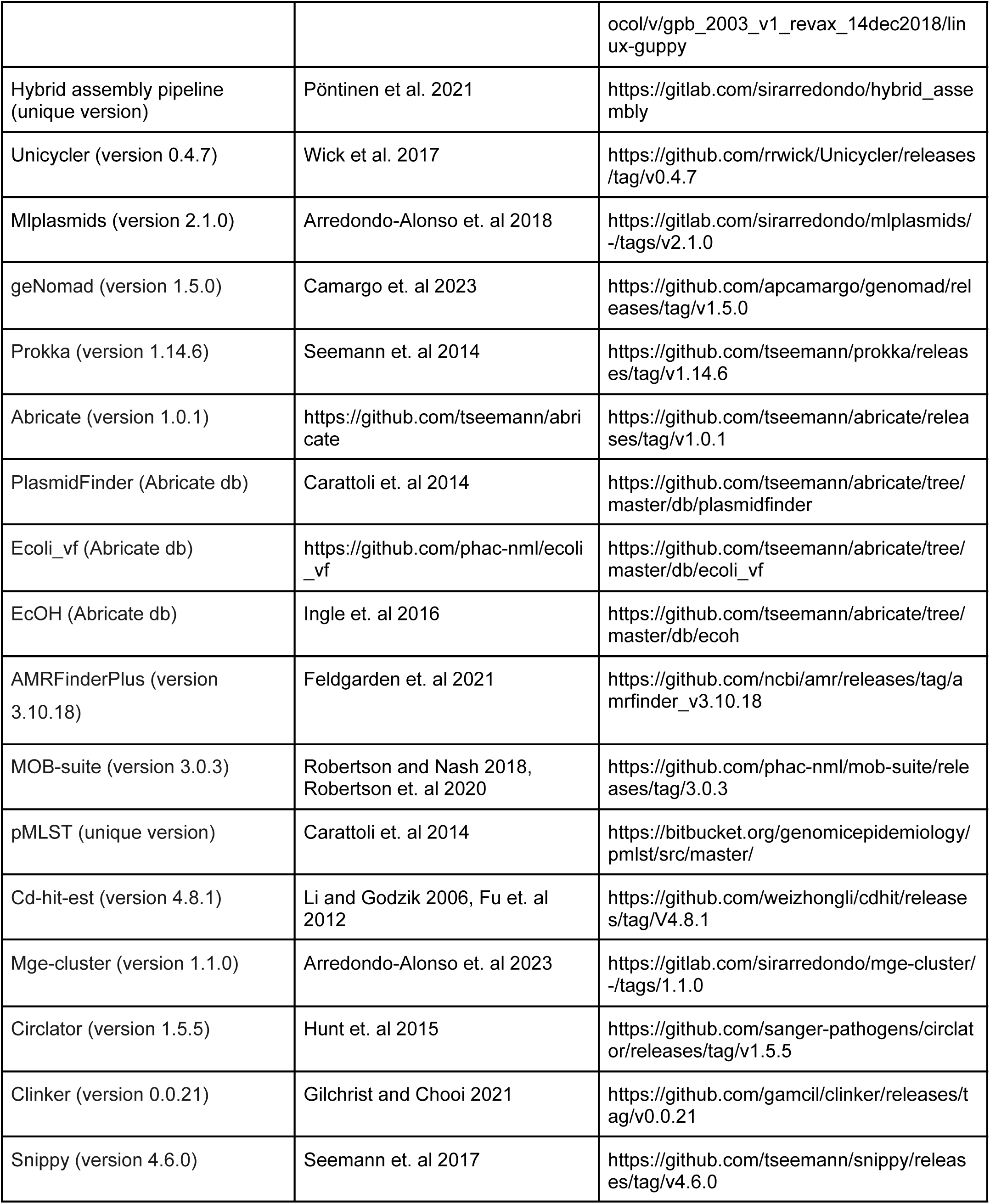
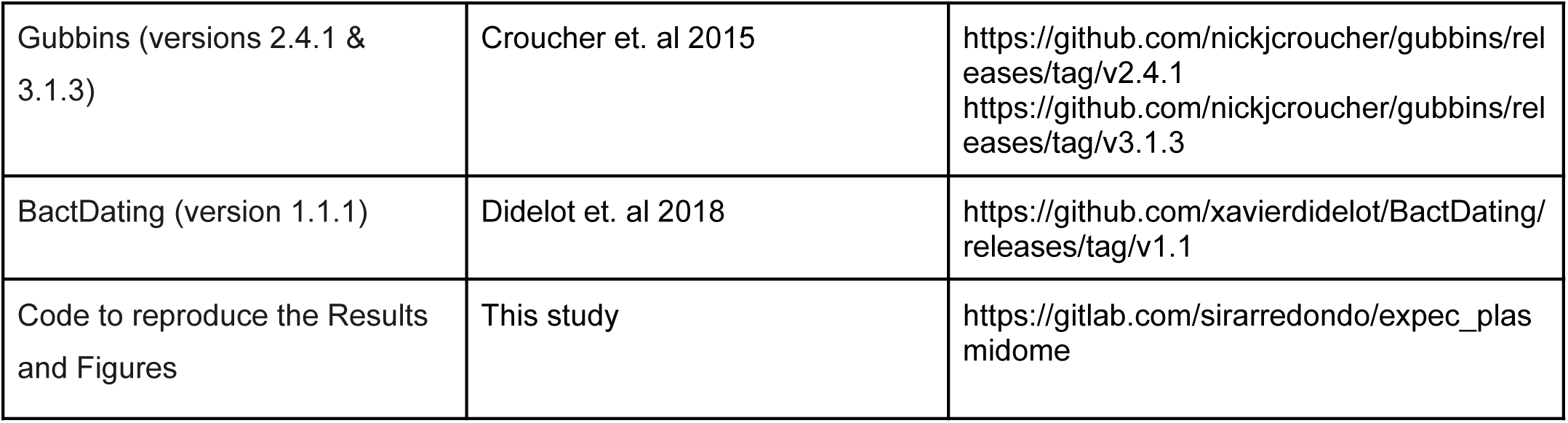

### Resource availability

#### Lead contact

Further information and requests for resources and reagents should be directed to and will be fulfilled by the Lead Contact, Jukka Corander.

#### Materials availability

Bacterial isolates and plasmids used in the bacteriocin experiments are available upon request to the Lead Contact, Jukka Corander.

#### Data and code availability

Short-read sequencing data have been deposited on ENA (Bioproject accession PRJEB32059). Long-read sequencing data and hybrid assemblies have been deposited on ENA (Bioproject accessions PRJEB45354 and PRJEB57633) and are publicly available as of the date of publication. Plasmid sequences derived from the hybrid assemblies are also publicly available at Figshare. DOIs listed in the key resources table.

The original code used to reproduce the analyses, figures and results presented in the manuscript is publicly available at https://gitlab.com/sirarredondo/expec_plasmidome

Any additional information required to reanalyze the data reported in this paper is available from the Lead Contact (Jukka Corander) upon request.

### Method details

#### Isolate selection for long-read sequencing

To resolve the accessory genome and mobile genetic elements (MGEs) of the extensive Norwegian Surveillance System for Resistant Microbes (NORM) ExPEC population ^11^, we selected 2,045 isolates (2,045/3,254, 62.85%) for long-read Oxford Nanopore Technologies (ONT) sequencing and, subsequently, for hybrid assembly. This selection was performed following a two-step strategy. First, to ensure that the resulting genomes represented the accessory genome diversity inherent in the NORM collection, 1,085 isolates (1,085/2,045, 53.06%) were selected regardless of their clonal complex using an unbiased statistical approach specifically designed for selecting representative isolates for long-read sequencing ^31^ (Fig. S1A). Briefly, we considered the presence/absence matrix of the orthologous genes computed by Panaroo (version 1.2.3) ^74^ on the 3,254 Illumina assemblies ^11^ and color the isolates based on their previously described PopPUNK (version 2.0.2) groups ^11,75^. From these, we used the k-means algorithm to select 1,085 isolates on the dimensionally reduced matrix computed by t-sne (perplexity = 30) (R stats package Rtsne, version 0.15) considering Jaccard distances as input.

Second, we focused on the four major ExPEC clones (ST73, ST95, ST131 and ST69) and sequenced all their remaining isolates not selected within the first step (960/2,045, 46.94%). This permitted us to directly estimate the prevalence of particular plasmid types among the four major ExPEC clones.

#### DNA isolation and ONT sequencing

Long-read sequencing of the 2,045 isolates was performed using a high-throughput multiplexing approach based on ONT reads ^31^. All the isolates were separately grown on MacConkey agar No. 3 (Oxoid Ltd., Thermo Fisher Scientific Inc., Waltham, MA, USA) at 37° overnight. Individual colonies were picked for overnight growth at 37° in 1.6mL LB (Miller) broth (BD, Franklin Lakes,NJ, USA) each. Genomic high-molecular-weight DNA was extracted from cell pellets using MagAttract R HMW DNA Kit (Qiagen, Hilden, Germany) to a final elution volume of 100 μL. Output concentration and DNA integrity were measured using NanoDropOne spectrophotometer (Thermo Scientific) and the Qubit dsDNA HS assay kit (Thermo Fisher Scientific) on a CLARIOstar microplate reader (BMG Labtech, Ortenberg, Germany). The samples were then adjusted to 400 ng for long-read sequencing. The ONT libraries were prepared using SQK-LSK109(-XL) or SQK-NBD110-96 barcoding kit for 24- and 96-barcoding runs, respectively, and 40 fmol per sample was loaded onto FLO-MIN106 flow cells. Sequencing was run for 72 hours on GridION (Oxford Nanopore Technologies, Oxford, UK) using MinKNOW Core versions 3.6.0 to 4.2.5. Basecalling and demultiplexing were performed using Guppy versions 3.2.8 to 4.3.4, with fast basecalling for the 24-barcoding and high-accuracy basecalling for the 96-barcoding runs.

#### Hybrid assemblies

ONT long reads were combined with the existing short-read data ^11^ using a previously published hybrid assembly pipeline ^76^ publicly available at https://gitlab.com/sirarredondo/hybrid_assembly. This pipeline is largely based on Unicycler (version 0.4.7) and was designed to automate the generation of (near-)complete genomes. In total, we could obtain a hybrid assembly for 1,999 genomes (1,999/2,045, 97.75%). The information derived from Unicycler regarding the length, circularity and depth of each contig was extracted and considered for the downstream analyses. The hybrid assemblies were split into individual contigs and circlator (version 1.5.5) ^77^ was used with the command fixstart to rotate the starting position of each contig.

Each contig was classified either as chromosome-derived, plasmid-derived or virus-derived (bacteriophage) considering both the predictions of mlplasmids (species ‘Escherichia coli’) (version 2.1.0) and geNomad (command end-to-end) (version 1.5.0) ^78^. First, contigs were labeled as chromosomal if they either had a size larger than 500 kbp or a mlplasmids chromosome probability higher than 0.7 and geNomad chromosome score higher than 0.7. Second, contigs were labeled as plasmids if they had a minimum mlplasmids plasmid probability of 0.3 and geNomad plasmid score of 0.7. These settings were used to compensate for the number of false negative predictions reported previously for mlplasmids using the *E. coli* model ^79^. Third, contigs not meeting any of the previous criteria were classified as virus-derived if they had a minimum geNomad virus score of 0.7. Fourth, the remaining contigs were considered as unclassified.

Prokka (version 1.14.6) ^80^ using the genus *Escherichia* and species *coli* was considered to annotate each of the contigs resulting from the hybrid assemblies. Abricate (version 1.0.1) (https://github.com/tseemann/abricate) coupled with the databases of PlasmidFinder ^81^, ecoli_vf (https://github.com/phac-nml/ecoli_vf), and EcOH ^82^ was used to assign the presence (minimum identity and coverage of 80%) of plasmid replicon sequences, *E. coli* known virulence factors (including colicin genes) and *E. coli* O-H serotype predictions respectively. AMRFinderPlus (version 3.10.18) ^83^ using the flag --plus and *Escherichia* as organism was also considered to identify antibiotic resistance, stress-response and virulence genes. The presence of plasmid replicon sequences was assessed using PlasmidFinder ^81^, while relaxases, mate-pair formation types and the predicted mobility of the contig was computed using the module mob_typer (version 3.0.3) of MOB-suite ^84,85^. The latter tool was used to extract the ‘primary_cluster_id’ which clusters sequences based on a fixed Mash distance threshold ^85^. Plasmid multi-locus sequence typing (pMLST) was computed using the IncF, IncA/C, IncHI1, IncHI2, IncI1, IncN and pbssb1-family schemes using the pmlst.py script available at https://bitbucket.org/genomicepidemiology/pmlst ^81^. For the plasmids with an IncF scheme annotation, we reported the FAB formula to contextualize them with previous literature.

### Typing the plasmid-derived sequences using a network approach based on mge-cluster

We removed the redundancy within the set of circular plasmid sequences using cd-hit-est (version 4.8.1) ^86,87^ using a sequence identity threshold of 0.99, alignment coverage of 0.9 and length difference cutoff of 0.9. Cd-hit-est computed a total of 2,560 clusters from which a single representative sequence was chosen.

The set of non-redundant plasmid sequences (n=2,560) was considered as input for performing the plasmid clustering based on mge-cluster (version 1.1.0) ^34^. This tool relies on two main parameters: perplexity and minimum cluster size. To avoid choosing an arbitrary parameter selection, we explored the hyperparameter space with: i) perplexity values ranging from 10 to 100 and ii) minimum cluster size values ranging from 10 to 50. By selecting 100 random combinations of these two parameters, we generated 100 distinct models with mge-cluster using the --create operational mode considering a unitig filtering variance of 0.01. For obtaining a robust plasmid assignment, we only considered plasmids with a minimum membership probability of belonging to a specific cluster (‘Standard Cluster’) of 0.8. The distinct mge-cluster models were combined into a single solution using a co-occurrence network approach. In this case, the network consisted of nodes belonging to plasmids (n=2,560) and edges to indirect connections with a weight ranging from 0 (plasmids are never part of the same cluster in the solutions) to 100 (plasmids are part of the same cluster in all solutions).

This network was scanned with the Louvain algorithm available in the R igraph package (version 1.2.6) to detect highly-connected communities within each non-singleton graph component, establishing a modularity threshold of 0.2. These communities were considered as plasmid types (referred as pTs) and are labeled using two digits representing their network component and their inferred community bin. For each pT, we discarded any sequences differing by two standard-deviations of the average plasmid length. This criteria was included to ensure that only plasmids with a similar size were part of the same pT.

To validate and contextualize the resulting pTs, the circular and non-redundant plasmid sequences of each pT were sketched (k=31,scaled=1000,noabund) with sourmash (version 4.8.2) ^36^ and a containment matrix computed using the command compare and the flag--containment. This matrix contains containment values corresponding to the fraction of a particular sketch found in a second sketch. The containment values were averaged within and between pTs. For each pT, we reported the most predominant (prevalence > 0.3) i) replicon found by PlasmidFinder, ii) pMLST scheme, iii) mobility (conjugative, mobilisable, non-mobilisable) as reported by MOB-suite, iv) cluster id (‘primary_cluster_id’) reported by MOB-suite, v) antibiotic resistance genes detected by AMRFinderPlus and vi) virulence genes detected by ecoli_vf (part of abricate). This information was summarized in Table S2.

The redundant sequences discarded by cd-hit (n=1,925) were assigned to the same pT as their representative sequence.

The results of the plasmid typing approach based on the co-occurrence network were compared to the 100 distinct solutions provided by mge-cluster using the adjusted Rand Index implemented in the R package mclust (version 5.4.7). The most similar solution provided by mge-cluster (perplexity = 62, minimum cluster size = 49) was used to visualize the clustering of the network approach by anchoring the sequences in a 2D embedding space. In addition, we used this solution in combination with the --existing operational mode to predict the position in the 2D embedding space of non-circular plasmid sequences (n=932) and the following well-studied ExPEC plasmids: CP000244 (pUTI89), CP007150 (pRS218), CU928146 (pECOS88), CU928148 (p1ESCUM), DQ381420 (pAPEC-O1-ColBM), EU330199 (pVM01), LQHK01000003 (pG150_1) and NC_022651 (pJJ1886_5). These sequences were assigned to the same pT as their closest (Euclidean distance) circular plasmid sequence. We only considered predictions differing by less than two standard-deviations of the average pT plasmid length.

### Plasmid synteny and phylogenetic analyses

For the pTs corresponding to large (>100 kbp) IncF plasmids pTs (1-1, 1-4, 2-1, 2-2, 2-3, 8-1, 8-2, 8-3, 9-1, 10-1, 13-1, 14-1)), we selected a plasmid sequence representative of the most common pMLST annotation and antibiotic resistance/virulence gene content of the cluster and with the earliest year of isolation (Table S2). The starting position of these sequences was changed to their IncFIB replicon using circlator (version 1.5.5) ^77^ with the command fixstart. Clinker (version 0.0.21) ^88^ with default settings was used to perform a gene synteny analysis and visualize which genomic blocks were shared among pTs, considering a minimum sequence identity of 0.8 (Fig. 2) and 0.99 (Fig. S6) to draw a link between genes.

We performed recombination-free phylogenies using the circular plasmid sequences from pTs 8-2&9-1&10-1 and 2-1&2-2&2-3. To create a core-genome alignment from these pairs of pTs, we used snippy (version 4.6.0) ^89^ which mapped the sequences from pTs &9-1&10-1 and 2-1&2-2&2-3 against the reference genomes 30348_1#293_2 (pT 10-1) and 30224_1#245_5 (pT 2-3), respectively. The resulting core genome alignment was polished using the module from snippy (snippy-clean_full_aln). Gubbins (version 3.1.3) ^90^ was used to compute a recombination-free phylogeny of the pTs using 100 bootstrap replicates, 50 algorithm iterations, and RAxML ^91^ as the application for model fitting and other default settings (GTRGAMMA as nucleotide substitution model).

### Dating the phylogenies of the four major ExPEC clones

The four major ExPEC clones were each mapped to a reference genome belonging to that lineage and recombination was removed using Gubbins v2.4.1 ^90^. Gubbins output was supplied to the R package BactDating v1.1.1 ^41^in three replicates and one with randomized tip dates. The R CaveDive package v0.1.1 was used to detect probable expansion events in the dated phylogeny ^42^. To report the dates of the distinct plasmid acquisition or introduction dates, we indicate in the text the lower and upper bounds of the confidence intervals (CI). For instance, pT 8-1 was acquired by ST131 from 1979 (95% CI 1971-1986) to 1984 (95% CI 1978-1989) (Table S2). This is reported in the text as acquired around ∼1981 (95% CI 1971-1989).

### Determination of bacteriocin activity

We produced four independent batches of microcin V by growing plasmid-harboring isolates 23-46, 27-20, 27-61 (ST95) and 28-33 (ST131 clade B) for 20 h at 37°C with aeration in 10 mL Lysogeny Broth (LB; LB Broth, Miller, Difco) with 0.2 mM 2,2’-bipyridyl (to limit iron availability; Sigma-Aldrich). The cultures were then centrifuged (4000 rpm, 10 min), filtered (0.2 μm polyethersulfone filter; VWR), and stored at 4°C. As controls, a plasmid-free ST131 clade B isolate 27-56 was treated in the same conditions, while isolate 27-61 was further treated without induction (no 2,2’-bipyridyl added).

To produce batches of colicin Ia, the four plasmid-harboring isolates mentioned above were grown in 10 mL LB overnight (15-16 h) at 37°C with aeration, centrifuged (4000 rpm, 10 min) and then the pellets were resuspended in 10 mL PBS. The suspensions were UV-irradiated (254 nm wavelength) with a dose of 36 mW/cm^2^/second for 10 seconds. 500 μL from these irradiated suspensions were inoculated into 9.5 mL LB and incubated 4 h at 37°C with aeration (covered in aluminum foil to prevent photoreactivation). The cultures were then centrifuged (4000 rpm, 10 min), filtered (0.2 μm polyethersulfone filter; VWR), and stored at 4°C. As controls, the plasmid-free isolate 27-56 was treated in the same conditions, while isolate 27-61 was further treated without induction (no UV-irradiation).

51 NORM isolates (Table S4), as well as *E. coli* MG1655, were grown in 1 mL LB in 96-deep-well plates overnight at 37°C with aeration. Indicator plates were prepared by inoculating 15 μL of each overnight culture into 15 mL LB top agar (LB with 0.75% agar; Sigma-Aldrich) with 0.1 mM 2,2’-bipyridyl (to limit iron availability and promote expression of the colicin receptor Cir). 10 μL of each bacteriocin and control batch were spotted on indicator plates, which were incubated overnight at 30°C after drying. These assays were performed three independent times. *E. coli* MG1655 was further exposed to 10-fold dilutions of the bacteriocin batches to verify the characteristic dose-dependent inhibition and exclude phage activity.

Most of the sensitive NORM isolates displayed sensitivity only against microcin V, while the six sensitive to colicin Ia were sensitive to both bacteriocins. Because the supernatants could contain both bacteriocins (since at least one seems to be spontaneously produced at low concentrations), and they have a common receptor in target cells, we added a further control to unequivocally test the sensitivity of these isolates to each of the bacteriocins. From genomic data, a few isolates were predicted to encode only one of the bacteriocins and we used two of those (isolates 31-16 and 31-17) for further controls. These two isolates were treated, with and without specific induction for each bacteriocin, as described above. The ST95 isolate 31-17 encodes only microcin V, and produces inhibition only when grown under iron-limiting conditions (Fig. 4B). Alternatively, ST131-B isolate 31-16 encodes only colicin Ia and always produces inhibition, albeit at stronger levels upon SOS induction (Fig. 4C). Combined, this shows that the spontaneously produced bactericion is colicin Ia. This isolate is sensitive against the supernatant of 31-17, verifying the loss of the microcin V genes (lack of the immunity genes). Each of these two supernatants was tested against the six isolates mentioned above, and both were inhibitory to all six (as displayed in Fig. S10, Table S4).

### Quantification and statistical analyses

#### Choosing long-read isolates spanning the genomic diversity present in the NORM study

To select isolates for long-read sequencing, we used the presence/absence matrix inferred by Panaroo (version 1.2.3) on the short-read genomes of the 3,254 isolates included in the NORM study. This matrix was embedded into 2D using the R package Rtsne (version 0.15). The k-means algorithm part of the R stats package (version 3.6.3) was used to allocate 1,085 centroids and the ratio of the between-cluster sum of squares (betweenss) and the total sum of squares (totss) was considered to assess if the chosen isolates represented the diversity present in the 2D embedding.

#### Hybrid assembly evaluation

The contiguity of the hybrid assemblies was given based on the N50 value reporting the sequence length of the shortest contig representing 50% of the total assembly length. This value was reported using exclusively contigs predicted as chromosome-derived which better reflects the contiguity status of a hybrid assembly as previously reported ^31^.

#### Comparing chromosome and plasmidome sizes of ExPEC clones

The total sequence length of contigs predicted as chromosome and plasmid (termed plasmidome) was compared among ExPEC clones using the pairwise.wilcox.test function in the R package stats (version 3.6.3) using the Benjamini-Hochberg method for accounting for multiple testing correction.

#### Differentiating vertical versus horizontal plasmid transmission in the phylogenies

The cophenetic.phylo function from R package ape (version 5.5) was used to extract pairwise distances between the pairs of tips of the ExPEC phylogenies and specific pT phylogenies. Pearson correlation implemented in the R package stats (version 3.6.3) was used to estimate the correlation between plasmid-based (recombination-free) and ExPEC phylogenies (core-genome based) distances. A positive correlation among the two distances was considered suggestive of vertical transmission of the plasmid in the ExPEC phylogeny indicating a shared evolutionary history. In contrast, the lack of correlation between the two distances was indicative of multiple acquisitions of the same pT in the ExPEC phylogeny.

#### Assessing the convergence of the dated phylogenies

The R package Bactdating (version 1.1.1) was used to date the phylogenies of the ST69, ST73, ST95 and ST131 clones. We considered three independent models (replicates) and a model with randomized dates to infer the significance of the given dates. Each model ran through Markov chain Monte Carlo (MCMC) chains of 100,000,000 generations sampled every 1000 states with a 10,000,000 burn-in using the Additive Relaxed Clock (ARC) model ^41^. The three replicate MCMC chains were deemed to have converged with Gelman diagnostic of approximately 1 for mu, sigma and alpha using the coda R package ^92^. We assessed whether the effective sample size (ESS) on the first replicate model was greater than 200 using the effectiveSize function of the coda R package ^92^ and determined that the true dates model was better than the randomized dates model.

## Acknowledgements

Collaborators forming The Norwegian *E. coli* BSI Study Group: Nina Handal (Akershus University Hospital), Nils Olav Hermansen (Oslo University Hospital, Ullevål), Anita Kanestrøm (Østfold Hospital), Hege Elisabeth Larsen (Nordland Hospital), Paul Christoffer Lindemann (Haukeland University Hospital), Iren Høyland Löhr (Stavanger University Hospital), Åshild Marvik (Vestfold Hospital), Einar Nilsen (Molde Hospital and Ålesund Hospital), Marcela Pino (Oslo University Hospital, Rikshospitalet), Elisabeth Sirnes (Førde Hospital), Ståle Tofteland (Sørlandet Hospital), Kyriakos Zaragkoulias (Nord-Trøndelag Hospital Trust).

We acknowledge the support from the Genomic Support Centre Tromsø, UiT The Arctic University of Norway for Oxford Nanopore sequencing and François Cléon for excellent technical assistance.

The project was funded by the Trond Mohn Foundation (grant identifier TMS2019TMT04 to A.K.P., R.A.G., Ø.S., P.J.J., and J.C.). The presented work has received funding from the European Union’s Horizon 2020 research and innovation programme under the Marie Skłodowska-Curie Actions (grant No. 801,133 to S.A.-A. and A.K.P.) and has also been supported by the European Research Council (grant No. 742158 to J.C.).

## Author contributions

J.C, P.J, Ø.S designed and sought funding for the study. A.P performed bacteria culturing, DNA extraction and quality control for long-read sequencing. S.A-A performed the hybrid assemblies, and plasmid clustering, statistical analyses. R.G performed the computational work corresponding to the phylogenetic dating and clonal expansion assessment. J.A.G performed the experimental and functional assays on plasmid-encoded bacteriocins. J.C, P.J, Ø.S, S.A-A, A.P, R.G wrote the manuscript which was improved and revised by all authors.

## Supplementary Figures and Tables

**Figure S1.**
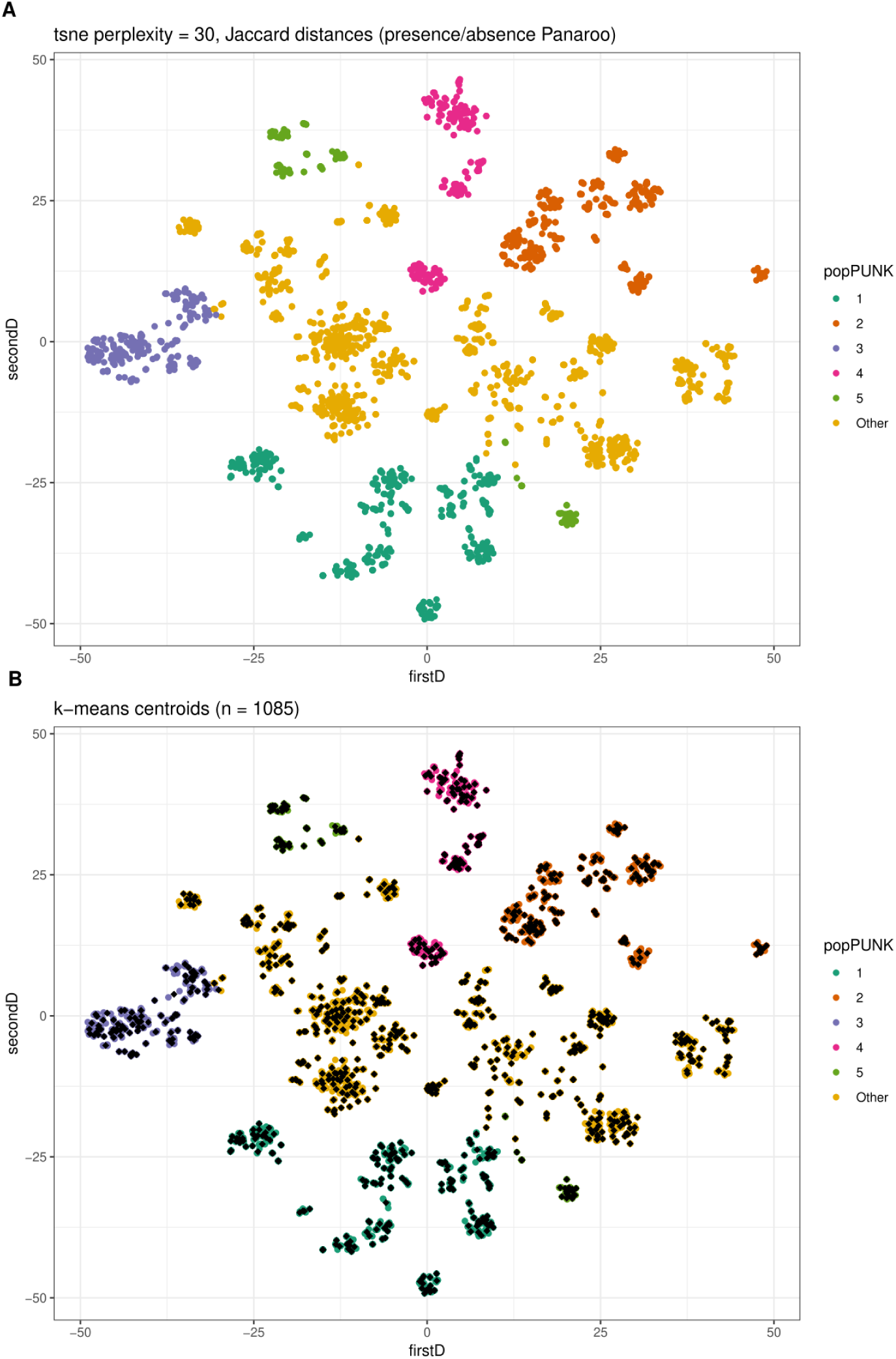
Selection of 1,085 ExPEC isolates for long-read sequencing based on their accessory genome diversity. A) Panaroo was used to infer the pangenome of the isolates included in the NORM collection (3,245) using short-read sequencing data. To represent the accessory genome diversity inherent in the collection, the presence/absence matrix of genes was embedded into 2D using t-sne (perplexity = 30) and isolates were coloured according to their PopPUNK group ^75^. B) To select 1,085 isolates for long-read sequencing, the k-means algorithm was used to locate 1,085 centroids in the 2D embedding. The isolate closest (Euclidean distance) to each centroid was selected for long-read sequencing.

**Figure S2.**
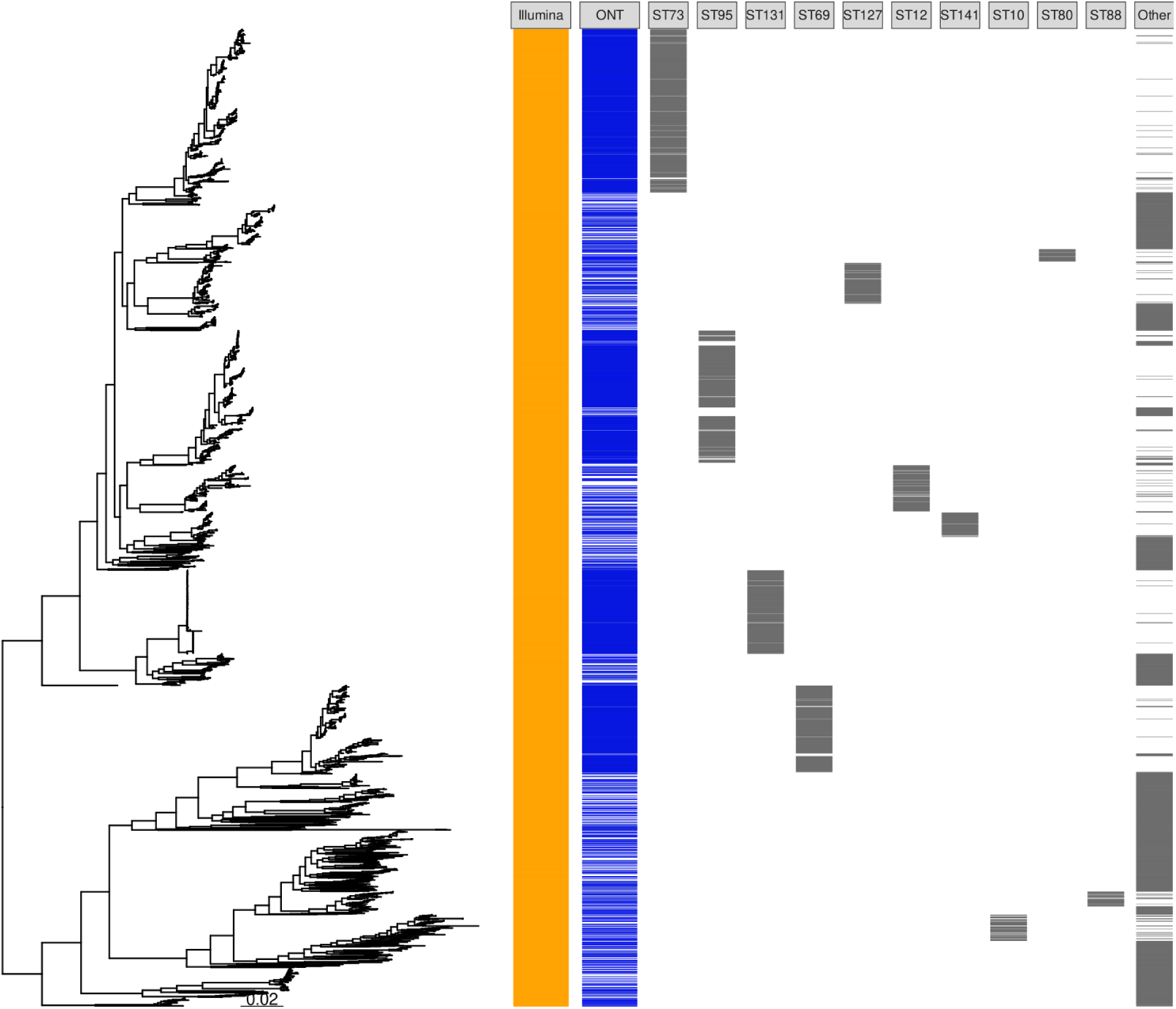
Distribution of the isolates selected for long-read sequencing (2,045 isolates, ONT column) in the ExPEC phylogeny (3,245 isolates). This selection included 1,085 isolates chosen based on their accessory genome diversity and 960 isolates belonging to ST69, ST73, ST95 and ST131 not included in the first selection.

**Figure S3.**
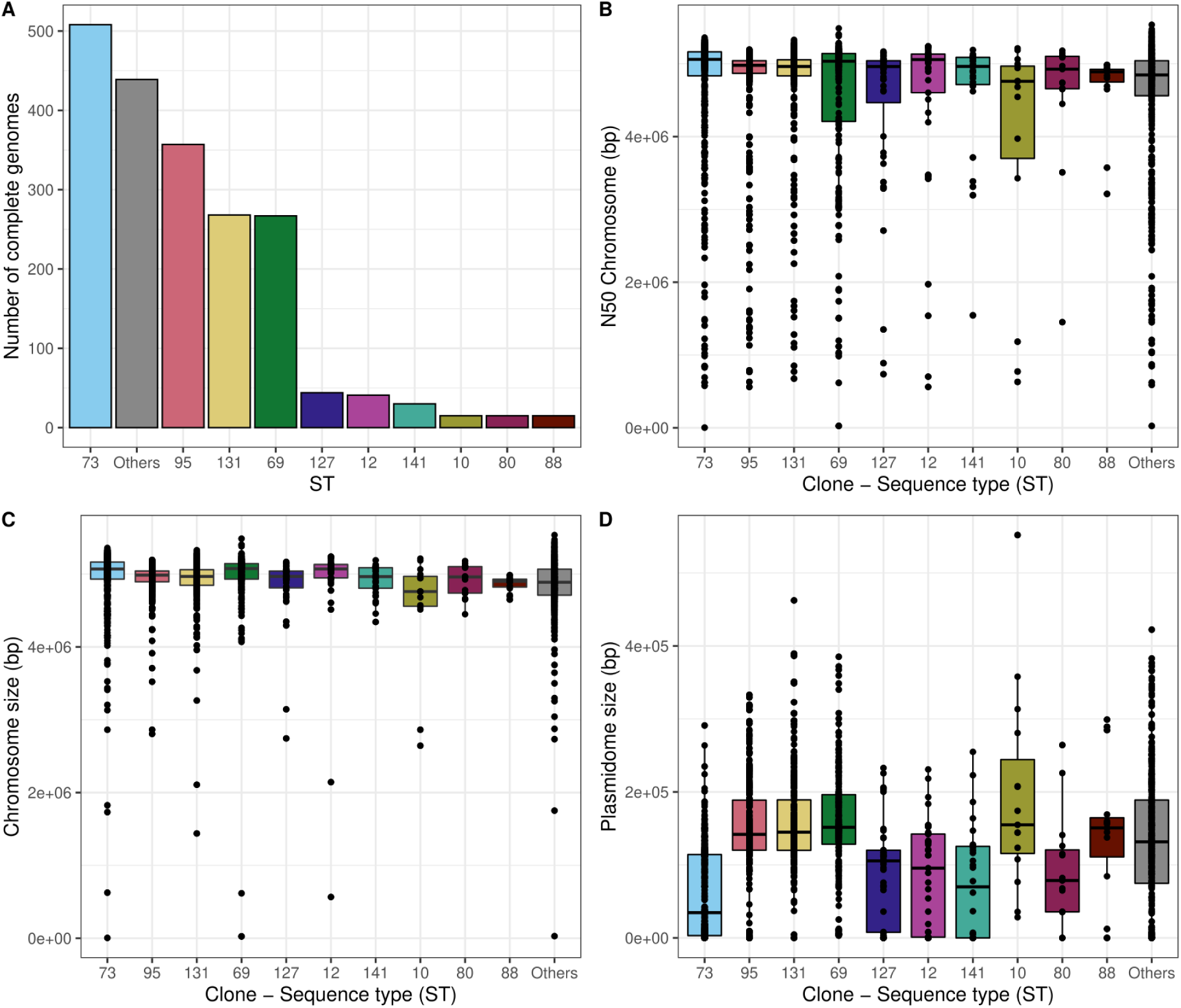
Genome statistics of the hybrid assemblies obtained for 1,999 ExPEC genomes. A) Barplot indicating the total number of assemblies available per ST. B) Boxplot of the N50 metric (in base-pairs) only considering contigs classified as chromosomal. C). Boxplot of the sum length (in base-pairs) considering contigs classified as chromosomal. D) Boxplot of the sum length (in base-pairs) of all contigs classified as plasmid (referred to as the plasmidome).

**Figure S4.**
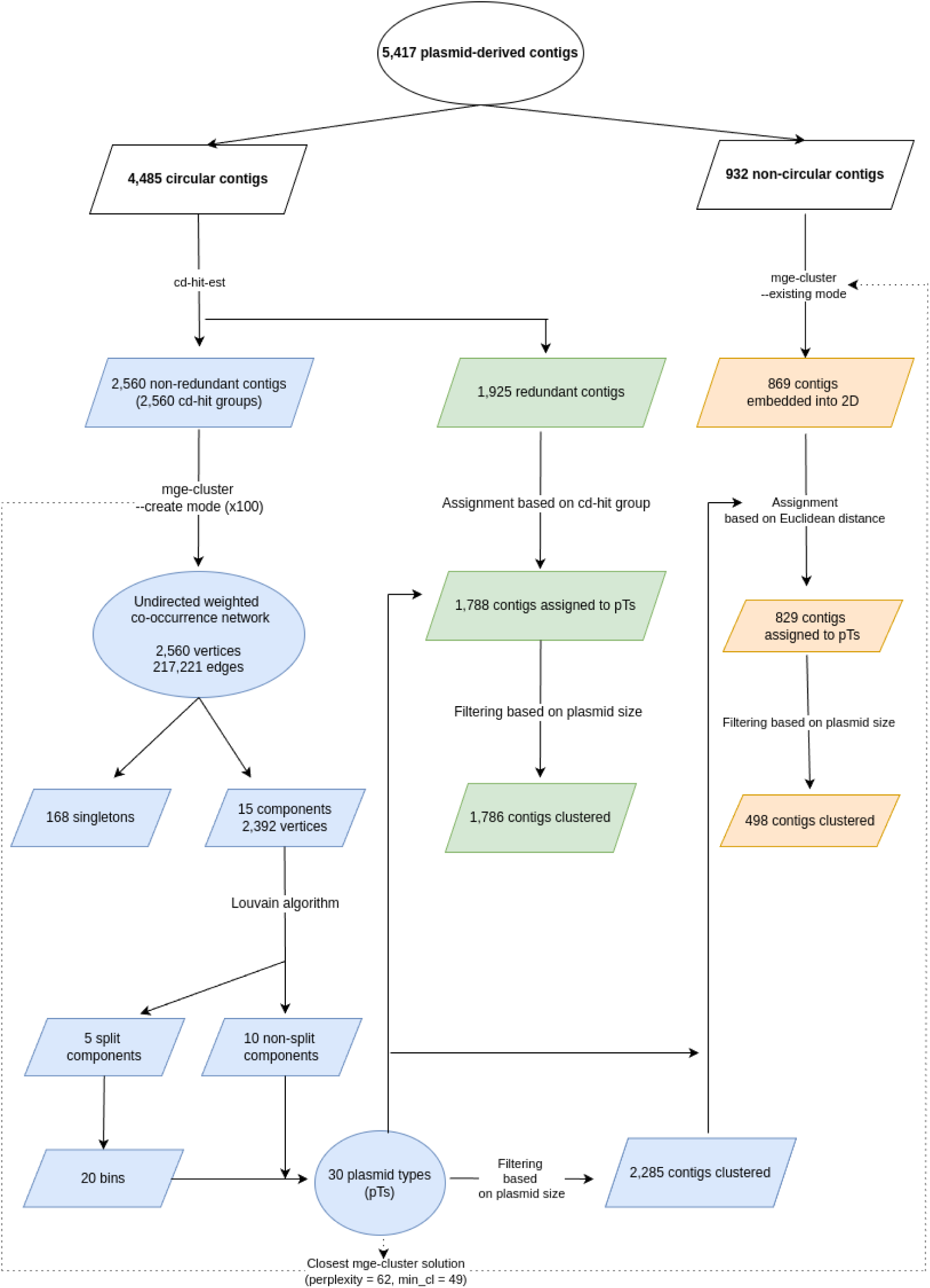
Workflow followed for performing the plasmid type (pT) assignment on the set of 5,417 plasmid contigs derived from the hybrid assemblies.

**Figure S5.**
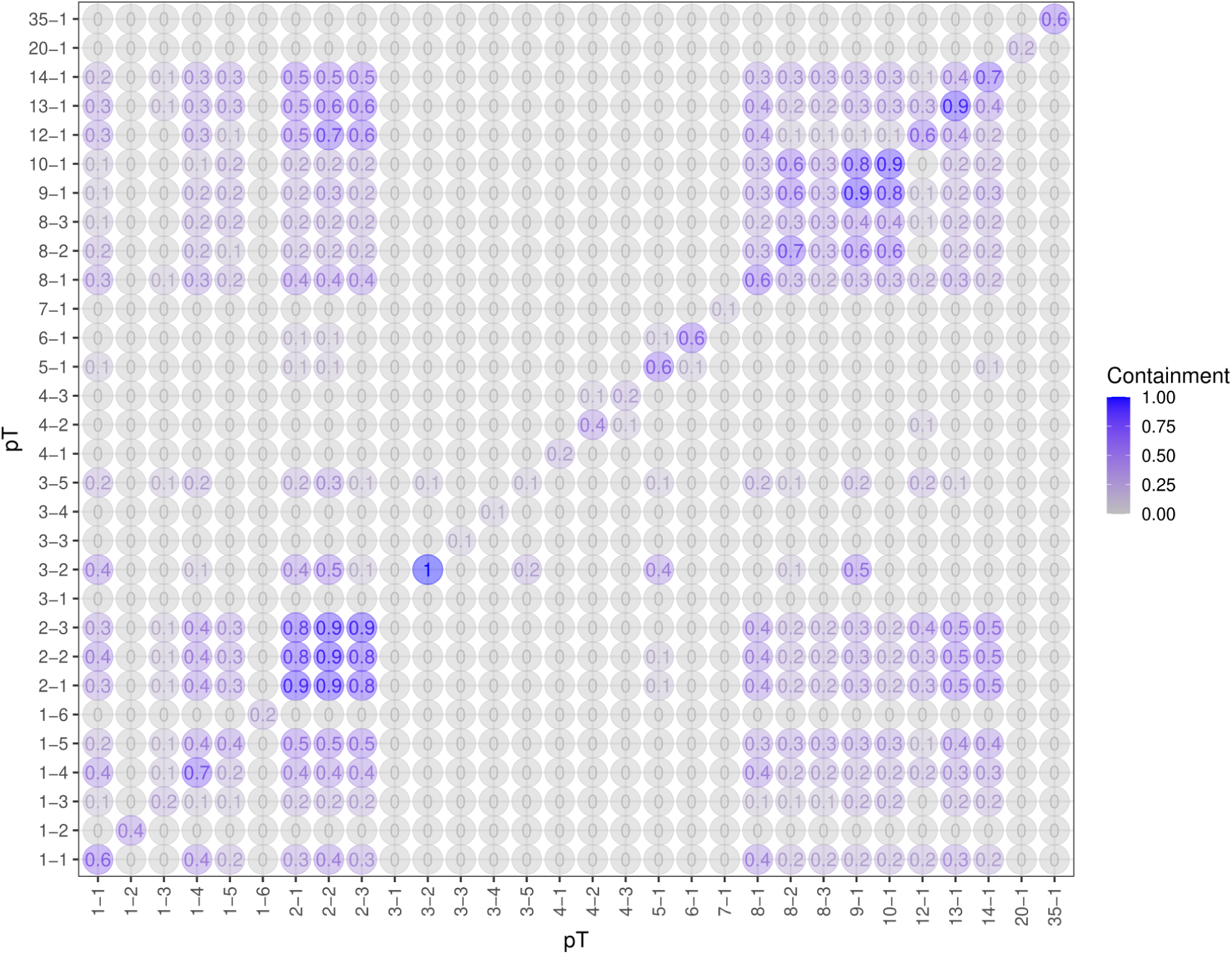
Sourmash containment analysis of the 30 plasmid types (pT) defined by mge-cluster. The containment value can range from 0 (pTs share no genomic region in common) to 1 (the entire genomic region of the pT indicated the y-axis is present in the pT indicated in the x-axis).

**Figure S6.**
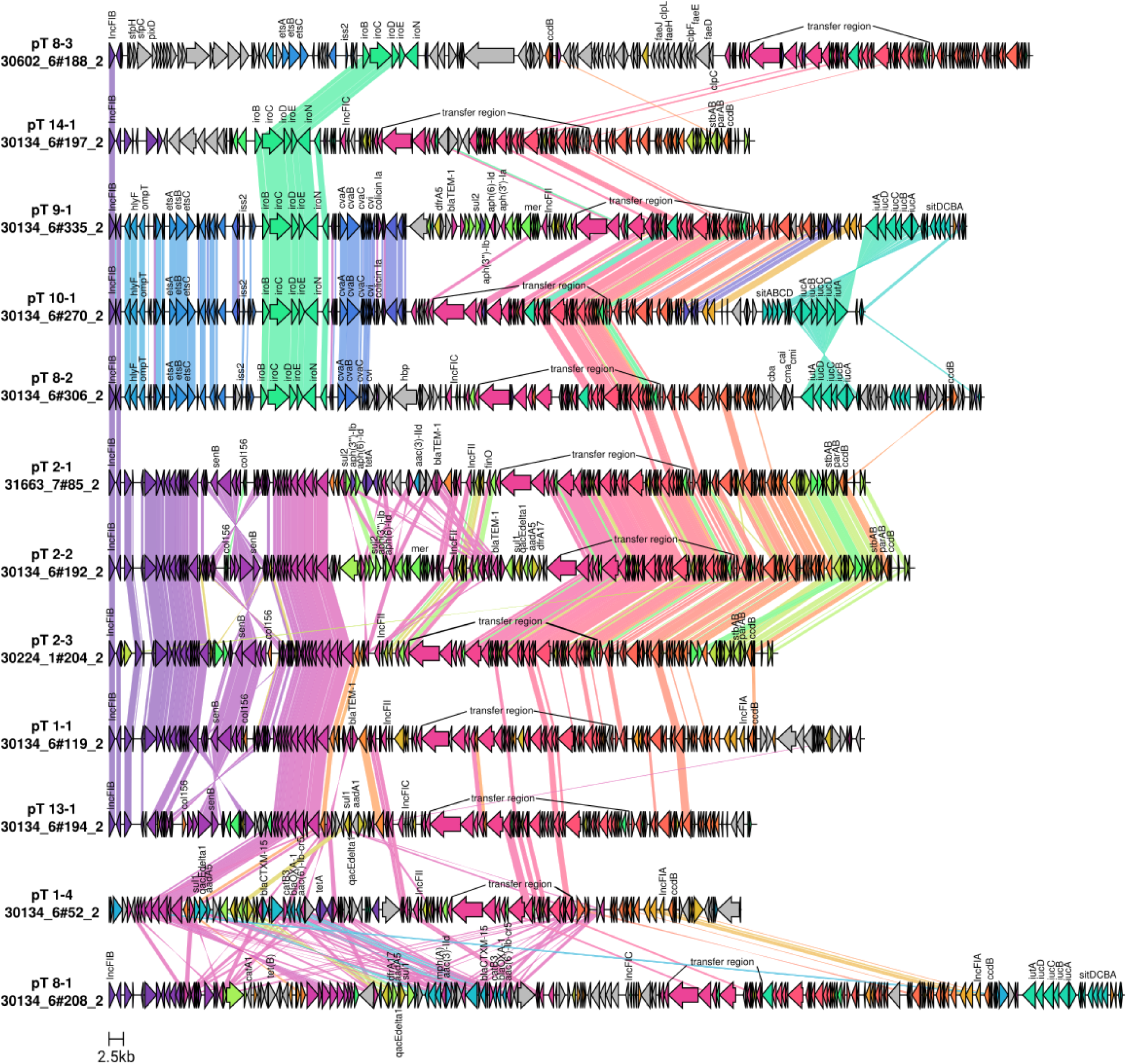
Clinker synteny visualization of the main large plasmid types (pT) considering a stringent minimum identity threshold of 0.99 to draw links between genes. Virulence and antibiotic resistance genes were identified using the EcOH database ^82^ (indexed in Abricate) and AMRFinderPlus ^83^ and their names curated according to well-studied ExPEC reference plasmids.

**Figure S7.**
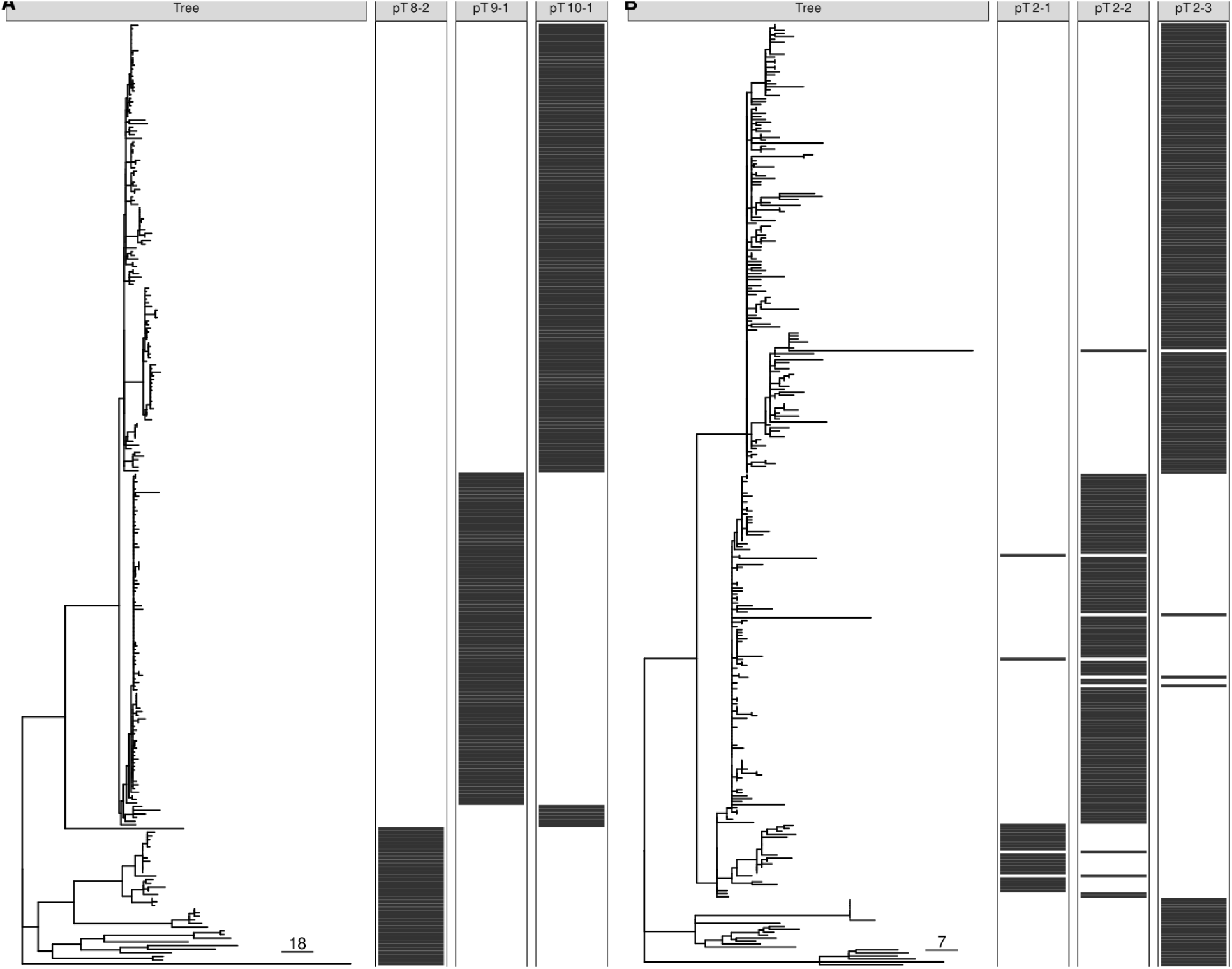
Recombination-free phylogeny based on plasmid sequences of pairs of plasmid types (pT) 8-2, 9-1, 10-1 (panel A) and 2-1, 2–2, 2-3 (panel B).

**Figure S8.**
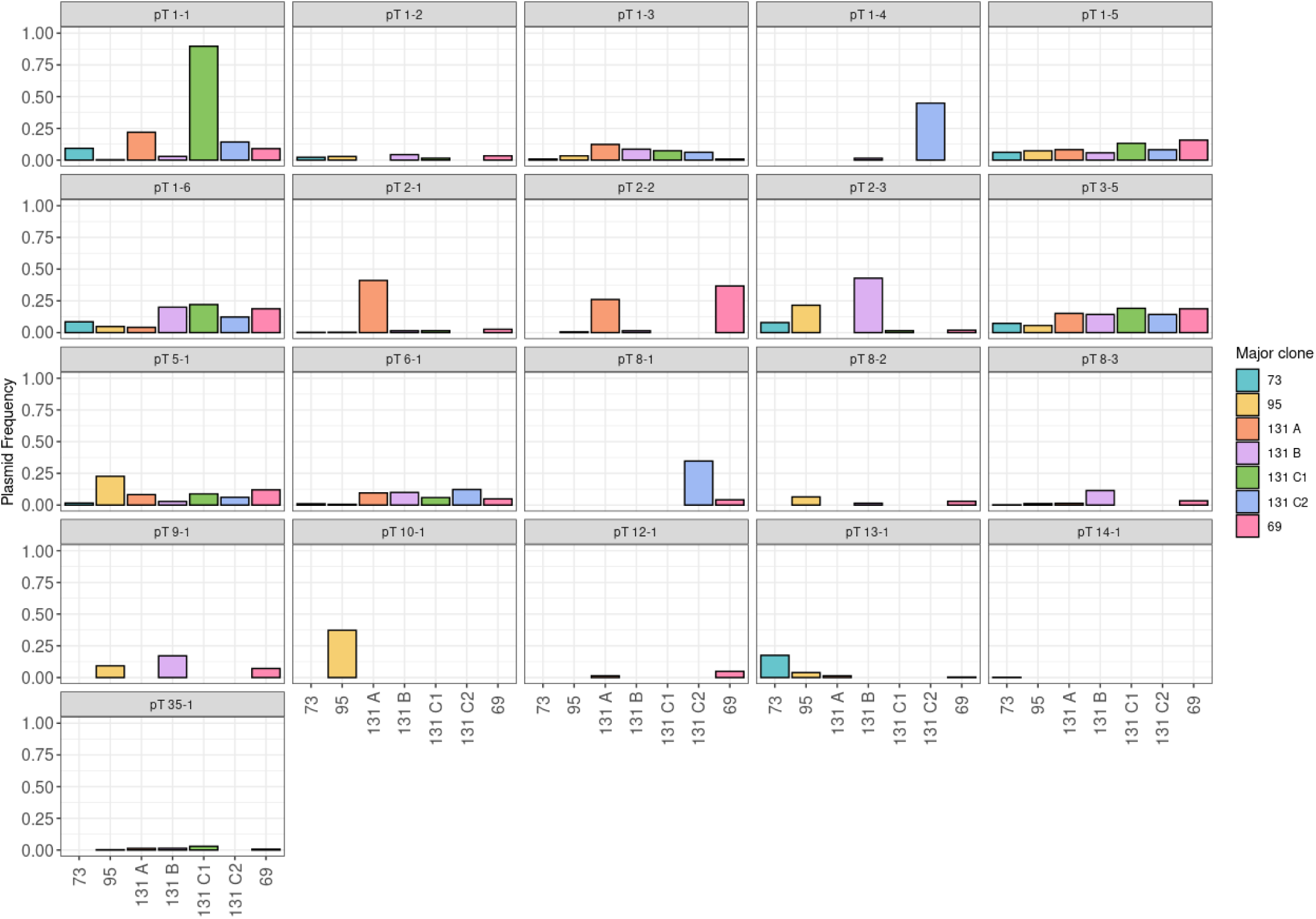
Plasmid type (pT) frequency among the four main ExPEC clones (ST69, ST73, ST95, ST131). In the case of ST131, we indicated the plasmid frequency according to the four clades present in the clone (A, B, C1, C2). Only pTs with an average size larger than 10 kb are represented.

**Figure S9.**
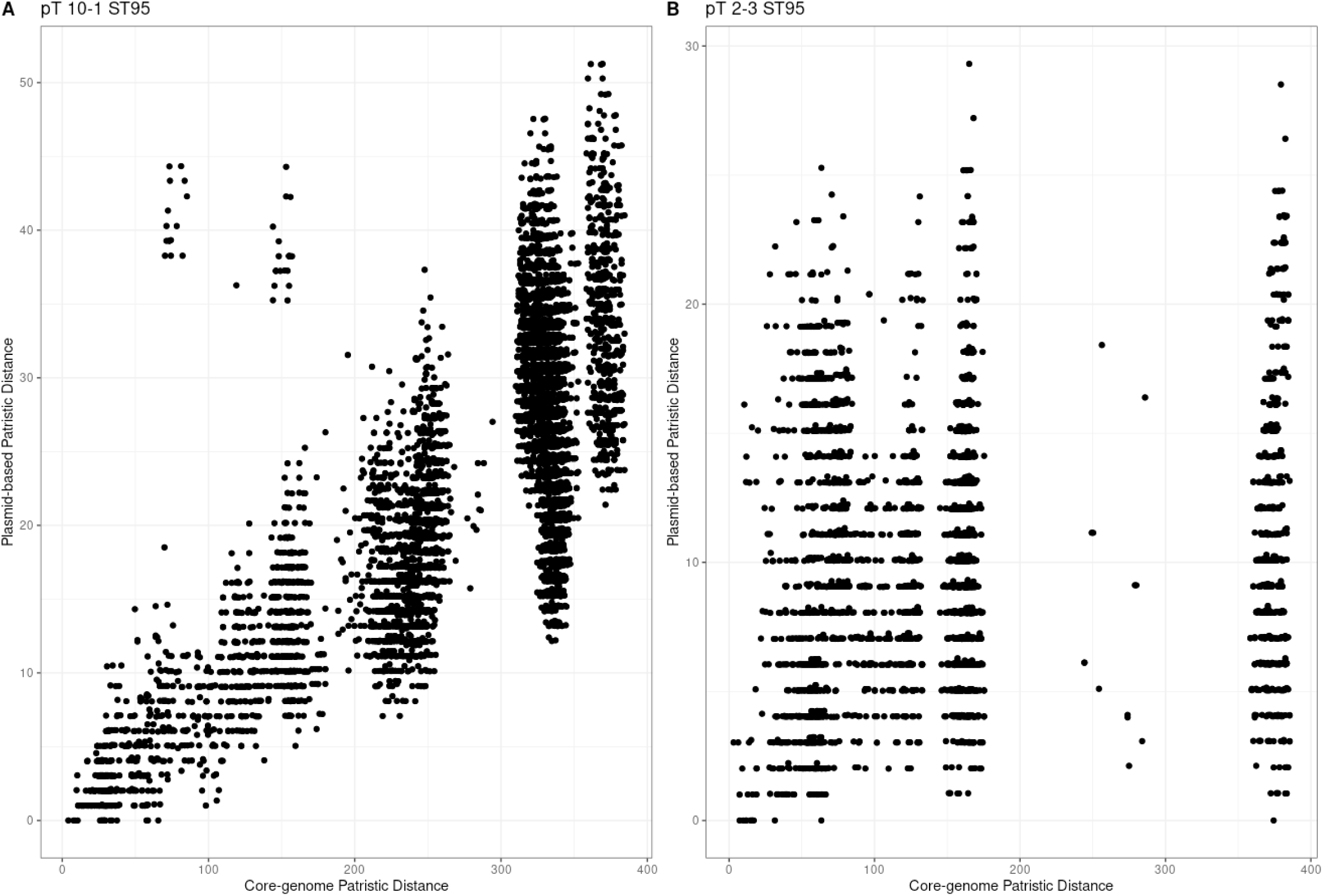
Plasmid-based versus core-genome patristic distances in the main plasmid types (pT) present in ST95. A) A positive correlation between the two distances was suggestive of vertical transmission of the pT 10-1 in the ST95 clone. B) The lack of correlation among the two distances suggested that pT 2-3 was introduced multiple times in the ST95 clone.

**Figure S10.**
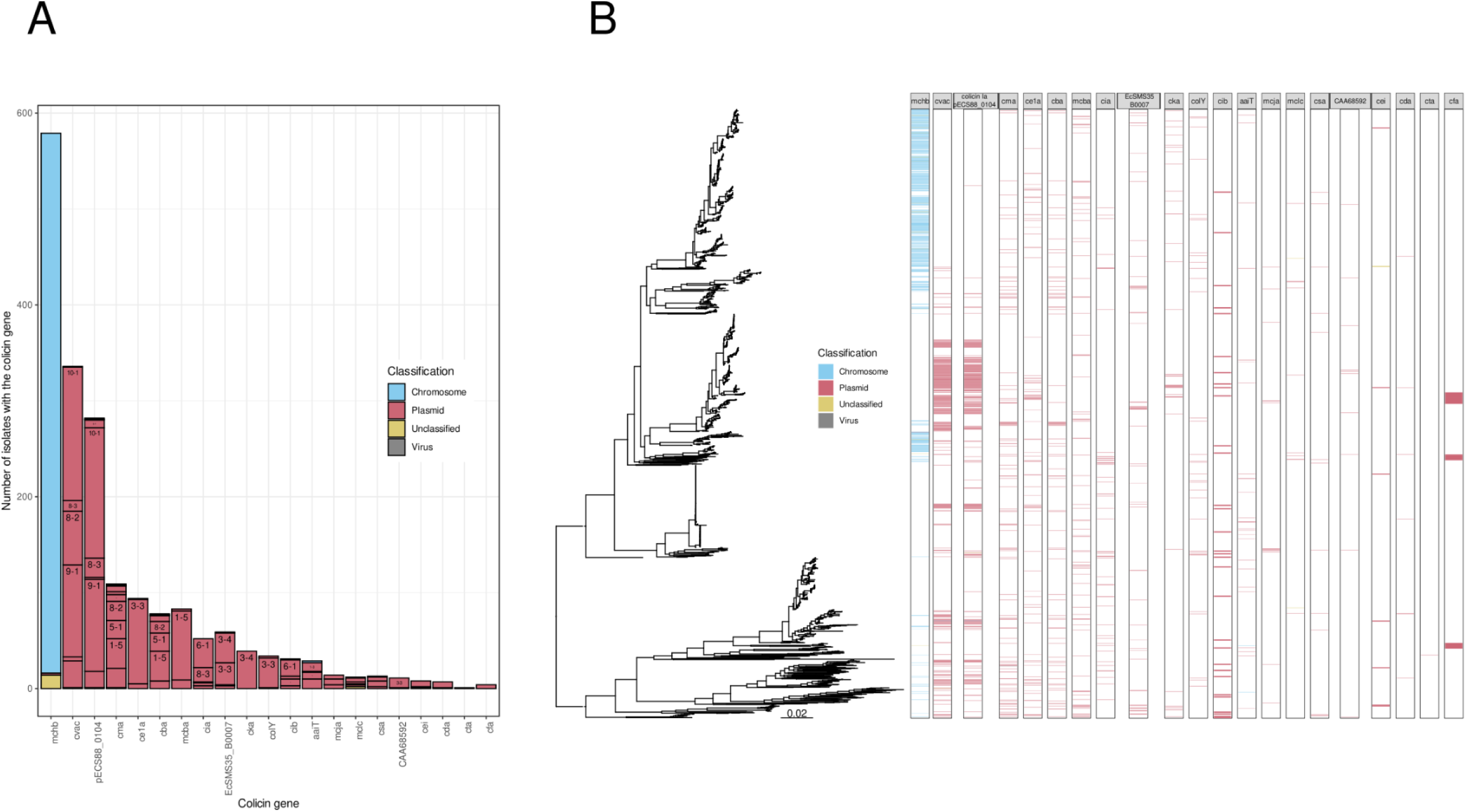
Distribution and classification of colicin genes in the ExPEC population. A) Stacked barplot showing the preferred genomic location of colicin genes: chromosome plasmid, virus (bacteriophage) or unclassified (unknown). In the case of plasmid location, we have indicated the particular plasmid type (pT) associated. B) Distribution of the colicin genes in the ExPEC phylogeny (as shown in Figure 1A). Each colicin was coloured according if the gene was encoded in the chromosome, plasmid, virus (bacteriophage) or an unclassified contig

**Figure S11.**
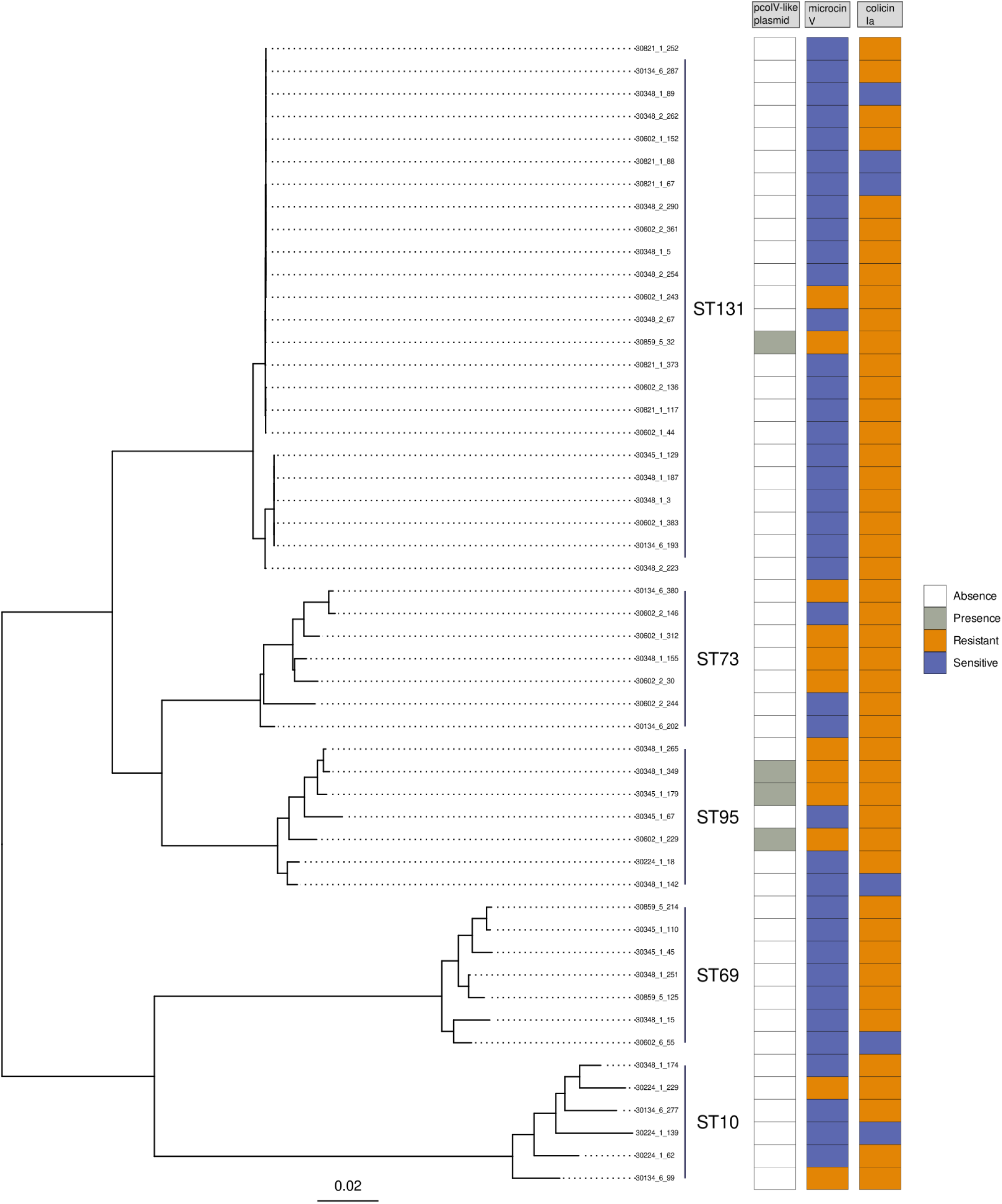
Bacteriocin susceptibility of NORM isolates. Pruned phylogeny of the 51 *E. coli* NORM isolates tested experimentally for bacteriocin susceptibility (right). Susceptibility to microcin V and colicin Ia is individually indicated in the heatmap in blue, while yellow indicates bacteriocin insensitivity (no effect detected). Isolates harboring the archetypical pcolV-like bacteriocinogenic plasmids are indicated in gray.

Supplementary Table S1. Report of the plasmid sequences (n=4,569) (4,071 circular and 498 non-circular) assigned to a particular plasmid type (pT) after quality control.

Each plasmid sequence is uniquely identified based on the ‘Plasmid_ID’ column and in the column ‘Cluster’ we provide its pT assignment. To contextualize each plasmid sequence with other typing schemes, we provide its PlasmidFinder, pmlst, MOB-Suite, antibiotic resistance (reported by AMRFinderPlus) and virulence (ecoli_vf database) annotations.

Supplementary Table S2. Plasmid type (pT) statistics and contextualization.

The number of sequences assigned to each type is reported after filtering out sequences initially predicted to belong to it, but differing by more than 2 standard-deviation of the average plasmid length reported. The reported predominant genes/categories are reported if they had a frequency of at least 0.3 and their exact frequency is reported in parenthesis. The name of the virulence genes reported by ecoli_vf (indexed in Abricate) was curated according to the annotation of well-known ExPEC plasmids.

Supplementary Table S3. Acquisition dates of the most prevalent plasmid types among the four major ExPEC clones.

Each acquisition is associated with a branch, but the exact time of introduction is unknown. For this reason, acquisitions are given as an interval date corresponding to the internal nodes flanking the branch. For each reported date, we provide the 95% confidence interval (CI) given by BactDating.

Supplementary Table S4. Isolates used in experiments. Internal collection numbers and NORM sequence identifiers (or alternative identifiers for lab strains) are given for each isolate.

Features identified in NORM genome sequences (plasmid pColV, bacteriocin genes, siderophore systems) are indicated as presence (+) or absence (-). Experimentally tested sensitivity to microcin V (col V) and colicin Ia (col Ia) is indicated as sensitivity (S) or insensitivity (I).

